# Deciphering the pangenome of the shellfish pathogen *Vibrio europaeus*: Evolutionary history and functional impact of core and accessory genes in aquaculture

**DOI:** 10.1101/2025.11.04.686615

**Authors:** Sergio Rodriguez, Diego Rey-Varela, Andrés Blanco-Hortas, Clara Martinez, Paulino Martínez, Marie-Agnès Travers, Juan L. Barja, Javier Dubert

## Abstract

*Vibrio europaeus* is an important pathogen in shellfish aquaculture, yet its genomic diversity and adaptive potential remain poorly understood. Here, we present the first comprehensive analysis of the *V. europaeus* pangenome, integrating genomic data from all strains available to the date sequenced specifically for this study. Those were isolated from different aquaculture facilities (shellfish hatcheries) associated to mass mollusk’s mortalities from different geographical locations, years and host species. Our findings revealed an open pangenome with the 61% of the genes associated to the accessory genome that contributes to environmental and host adaptations. Phylogenomic analyses of the core-genome (39% of the pangenome size) allowed to evaluate the evolutionary history and intraspecific diversity of *V. europaeus* and revealed that Spanish strains displayed a much lower genetic variability than French, Chilean or American strains, probably due to a monophyletic radiation event. Functional annotation of core and accessory genes revealed the key virulence factors of the species while it also disclosed that those are located mainly into the core genes. The high number of anti-phage defense systems encoded in the accessory genome explained almost all the variability of the species. The results provide important insights into the evolutionary history and ecological versatility of *V. europaeus*, with potential implications for diagnostics, epidemiological surveillance, and disease management strategies in aquaculture.

**Impact statement:** This study presents the first comprehensive pangenome analysis of *Vibrio europaeus*, an emergent pathogen responsible for severe economic losses in shellfish aquaculture, the second most important sector of global aquaculture. Here, we characterized for the first time the *V. europaeus* pangenome, integrating genomic data from all strains isolated to date, sequenced specifically for this study using NGS and/or third-generation (PacBio) technologies. This work achieved the most complete species pangenome to date and is among the first studies on aquaculture-related bacterial pathogens. Beyond a descriptive framework, the pangenome was critically examined to identify key traits, including virulence factors, secondary metabolite biosynthesis, and antimicrobial resistance genes, essential for host infection and adaptation. Moreover, the study of anti-phage defense systems was shown to account for much of the species’ genomic variability. The genomic resources and insights generated here substantially expand our understanding of *V. europaeus* biology and provide valuable information that can be applied for diagnostics, epidemiological surveillance, and sustainable management of this pathogen in aquaculture industry.

**Data summary:** All genome assemblies have been uploaded to the National Center for Biotechnology Information. The GenBank accession numbers for each of the 39 strains used in this study and detailed information can be found in Table S1. All bioinformatics tools used for comparative genomics have been listed in the Methods section including references, associated databases and analysis parameters.

## 1. INTRODUCTION

The term bacterial pangenome has been defined as the whole gene repertoire of a microbial species (McInerney, 2017). Advances in the study of the bacterial pangenome increase the understanding of the genomic adaptations of bacteria to the environment and contribute to the taxonomic and evolutionary research of the species (Liao, 2021; Fu, 2021; Simonsen 2022). The construction of a pangenome reflects indeed the diversity within a bacterial taxon and also enables the identification of common genes across strains or lineages, usually defined as the core genome. In opposition, the accessory genome would encompass the heterogeneity of genomes across strains or populations (McInerney, 2017). The core genome can be compared at various taxonomic levels providing the genomic basis of the species phylogeny (McInerney, 2017). Additionally, the accessory genome importance relies on the fact that enables bacterial adaptation to ecological niches, providing the variability underlying specific traits such as a virulence, defense systems against bacteriophages, antimicrobial resistance or production of secondary metabolites (Jackson, 2011; Vassallo, 2022). Pangenome analyses are an excellent approach to study pathogenic strains facilitating the study of its adaptation to the environment and hosts.

Bivalve aquaculture is considered the second most important activity within the aquaculture sector, only surpassed by the algae production (FAO 2024), however its expansion is constrained by the negative impact of bacterial diseases, causing great economic and animal losses to the industry. One of the most prevalent diseases is the vibriosis caused by different pathogenic *Vibrio* spp. (Prado, 2005; Dubert, 2017b). Though, vibrios are commonly considered as opportunistic pathogens, some species from this genus are adapted to infect a variety of hosts and bear virulence elements implied in different infection stages, such as bacterial adherence, invasion and survival in the host’s environment (Shapiro-Ilan, 2005; Le Roux, 2016; Parizadeh, 2018; Destoumieux-Garzón, 2020). Among *Vibrio* pathogens in shellfish, *V. europaeus* is an emergent species with great impact on bivalve aquaculture worldwide, affecting the most important mollusks species and infecting even different stages of development (i.e., broodstock, eggs, larvae, post-larvae and juveniles). *V. europaeus* has been identified in outbreaks on the main producer countries such as Spain, France, Chile or US (Prado, 2005; Mersni-Achour, 2014; Travers, 2014; Prado, 2015; Dubert, 2016d; Dubert, 2017c; Rojas, 2021).

Pan-genome analyses are key to obtain key insights into the species’ genomic diversity and adaptive potential, with implications for understanding its evolution and guiding applications such as secondary metabolite discovery and the development of preventive strategies. However, the pangenome studies about bacterial pathogens affecting bivalve aquaculture are very scarce (Dias, 2018). The aim of this study was to characterize for the first time the pangenome of the bivalve pathogen *V. europaeus* to gain knowledge about the genomics of this species. For this purpose, a collection of all *V. europaeus* strains isolated to the date was sequenced and used to construct the bacterial pangenome. Core and accessory genes were identified and different comparative analyses were performed to elucidate questions related to its evolution, intra-specific diversity, virulence, production of secondary metabolites and resistance.

## 2. MATERIAL AND METHODS

### 2.1. V. europaeus strains: DNA extraction, whole genome sequencing and assembly

All strains identified to date as *V. europaeus* (39 strains) were used for the pangenome reconstruction, including 36 *V. europaeus* strains sequenced in this study and three genomes retrieved from NCBI (strains 071316F, NPI-1 and CECT8426; Table 1). The 36 strains sequenced were obtained from 6 different hatcheries located between Spain (3), Chile (1) and France (2), and one environmental sample from US (Table 1). Two sequencing approaches were followed for genome assembly:

i. Short-read sequencing: 150 bp paired-end Illumina sequencing was performed in the *V. europaeus* strains (36 strains) for assembling at the contig level (Table 1). Strains were grown overnight in Trypto–Casein Soy agar supplemented with 2% (w/v) sodium chloride (TSA–2, Condalab) at 25°C. Subsequently, a single colony was picked and grown under the same bacterial culture conditions in broth (TSB-2) with vigorous shaking. DNA was extracted from an aliquot of the overnight culture (200 µl) using the DNeasy Blood & Tissue Kit (QIAGEN) following the manufacturer instructions. Quantity, quality and integrity of each DNA extraction was evaluated using NanoDrop One (Thermo Scientific), Qubit (Thermo Scientific) and by electrophoresis in a 1% (w/v) agarose gel. Genomic libraries and sequencing (HiSeq4000 sequencer, Illumina) were performed by the SNPsaurus company. Quality of Illumina reads was performed using Trimmomatic (Bolger, 2014) and paired-end short reads were assembled at contig level using SPAdes 3.15.4 (Prjibelski, 2020).
ii. Long-read sequencing: Four representative strains (the type strain CECT8136 and the isolates CECT8427, PP2-843 and EX1; Table 1) were long-read sequenced to achieve high resolution assemblies polished with Illumina reads from the previous section. For this, each bacterial strain was grown as described above and HMW DNA was extracted from 4 ml of an overnight culture using Qiagen Genomic-tips 100/G kit (QIAGEN). Quantity, quality, and integrity of HMW DNA extractions were evaluated as described above. CECT8136, CECT8427 and PP2-843 strains were sequenced using a PacBio Sequel II sequencer (PacBio) by SNPsaurus, while EX1 genome was sequenced in using MinION sequencer with the Rapid sequencing gDNA-barcoding kit (ONT). Assemblies from PacBio and ONT reads were performed using de novo long-read assembler Flye (Kolmogorov, 2019) and polished by Racon (Vaser, 2017). Finally, each assembly was polished using Illumina reads with Pilon (Walker, 2014). Thus, a total of six whole genome chromosome-level assemblies (CECT8136T, CECT8427, PP2-843 and EX1 sequenced in this study; and NPI-1 and CECT8426 retrieved from NCBI) were used in this study for pangenome analyses (Table 1).

**Table 1.**
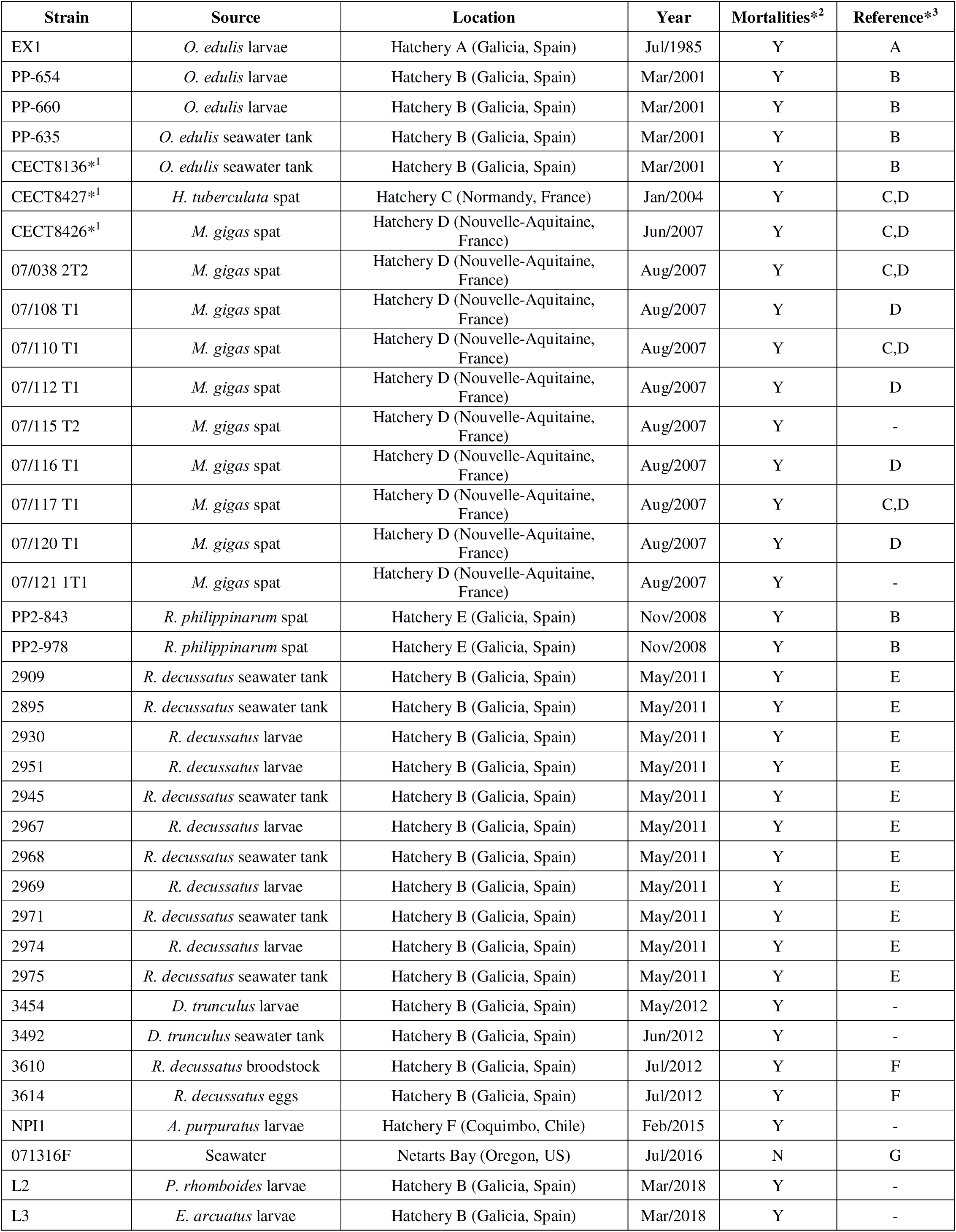
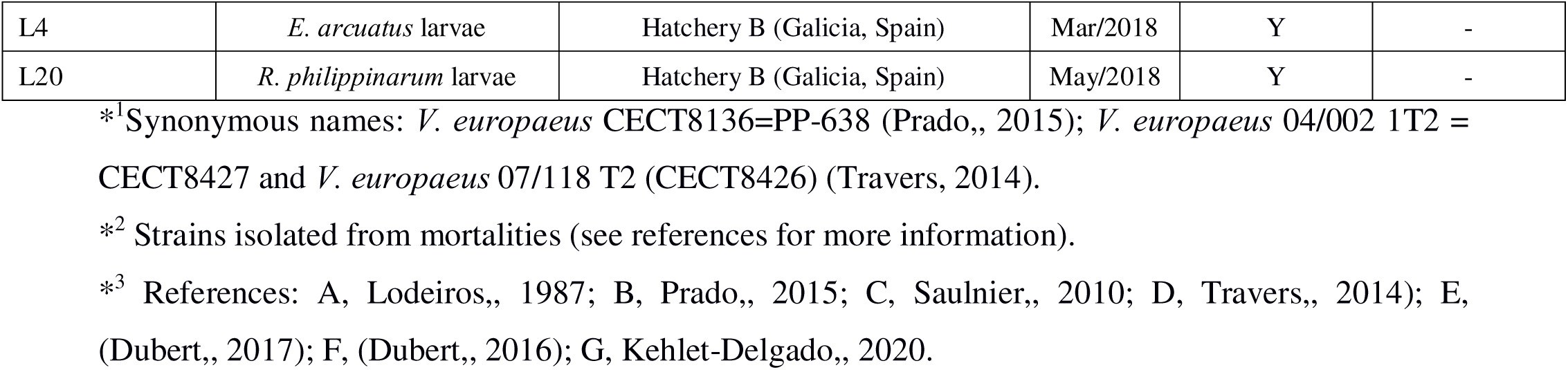
Main features of all *V. europaeus* strains (39 strains) identified to date.

### 2.2. Genome annotation and pangenome construction

Each genome assembly (39 genomes; Table 1) was annotated by Prokka 1.14.4 (Seemann 2014). The gff3 file was used to construct the *V. europeus* pangenome using Roary 3.13.0 under default parameters (Page, 2015). The core genome includes those genes that were present in the 100% of the *V. europaeus* genomes, while the accessory genome was further classified as shell core, soft core and cloud genome if a gene was present in the 95-99%, 15-95% or 0-15% of the genomes, respectively. Genes classified into core, shell core, soft core or cloud fractions were plotted with ggplot2 3.3.6 R package (Wickham 2016).

The micropan R package (Snipen and Liland 2015) was used to determine the *V. europaeus* pangenome aperture by the calculation of the Heaps alpha value setting 1000 permutations and results were plotted as described above.

Gene sequences from each pangenome fraction (i.e., core, shell core, soft and cloud) were retrieved from the Roary’s pan_genome_reference file using SeqKit 2.1.0 (Shen, 2016). Subsequently, each gene was functionally annotated using reCOGnizer 1.7.0 (Sequeira, 2022) to obtain the distribution of Cluster of Orthologous Group (COG) categories within the *V. europaeus* pangenome. The percentage of genes assigned to the different COG categories per pangenome fraction was calculated and the resulting matrix was plotted in a heatmap using the ComplexHeatmap 2.11.1 R package (Gu, 2022).

Identification of tRNAs and rRNAs from each genome assembly was done using tRNAscan-SE 2.0.1 (Chan, 2021) and barrnap 0.9 (https://github.com/tseemann/barrnap) respectively.

### 2.3. Phylogenomic comparisons: phylogenetic tree, SNPs identification and Average Nucleotide Identity (ANI) calculation

Genes from the core fraction of each genome assembly (39 genomes) were aligned by Roary 3.13.0 using MAFFT. This alignment was used to build a phylogenomic tree with IQ-TREE (Nguyen, 2015) based on the maximum likelihood (ML) algorithm with the UNREST model by bootstrapping over 1000 replications and the phylogenomic tree was visualized in iTOL v6.6 (Letunic and Bork 2021).

Single nucleotide polymorphisms (SNPs) were identified across the core genes alignment created with MAFFT using an *ad hoc* Phyton script (available in https://github.com/sergio-c-r/core-snps/blob/main/snp_dif.py). Average nucleotide identity (ANI) among the different *V. europaeus* genome assemblies was calculated by PYANI 0.2.11 (Pritchard, 2016) with default parameters and plotted in a heatmap as described in 2.2.

A dendrogram of the accessory gene profiles was constructed though hierarchical clustering using R, based on the presence/absence matrix generated by Roary. To illustrate the presence/absence genes profiles, the results were plotted as a heatmap using the R package ComplexHeatmap 2.11.1, including the dendrogram. Finally, a principal component analysis (PCA) was performed based on the presence/absence matrix of pangenome genes by R.

### 2.4. Experimental infections

A total of 38 *V. europaeus* strains (71316F was not available in our bacterial collection; Table 1) were tested in virulence challenges using Manila clam (*Ruditapes philippinarum*) juveniles (14±1 mm). Virulence challenges were performed following the infection protocol described by Martínez, 2022, *Vibrio breoganii* C5.5 was used as negative control. Briefly, bacterial strains were grown overnight, and bacterial suspensions were made in sterile sea water (SSW) adjusted to OD_600_=1 and confirmed by decimal dilution series onto TSA–2 plates (∼10^8^ CFU ml^−1^). Experimental challenges included two steps:

i. Infection: tanks were filled with filtered seawater (FSW; 0.22 μM Nalgene Rapid–Flow, Thermo Scientific) containing a bacterial suspension adjusted to a final concentration of 10^7^ CFU ml^−1^. Subsequently, 15 Manila clam juveniles were added to each tank and kept for 24 h at room temperature (RT=∼20°C) for active bacteria filtration. Experimental challenges were performed in triplicate.
ii. Post–infection: challenged juveniles were removed from the infection tanks and maintained for 8 h at RT to allow the internalization of the bacteria within the pallial cavity. Then, juveniles were transferred to new tanks filled with 200 ml FSW and maintained at RT with aeration. Mortalities were monitored at 8 h, 20 h, 32 h, 44 h, 56 h, 68 h and 80 h post–infection and clams were immediately removed when the valves were open (dead juveniles), or siphons were not retracted following stimulation (moribund juveniles). Results were recorded as a percentage of survival. FSW was renewed once per day (or if it was turbid).

### 2.5. In-silico identification of the virulence genes

Genes related to virulence were identified in each *V. europaeus* genome (39 genomes) using the VFDB database (Liu, 2021). An additional set of curated *ad oc* database proteins whose role in virulence experimentally demonstrated was included in the analyses by BLASTp comparisons. The resulting presence-absence matrix was used to plot a heatmap as described in 2.2. Subsequently, a principal component analysis (PCA) was performed using the prcomp fuction available in R base and plotted with ggplot2 to compare the virulence genes belonging to the non-core fractions among the different *V. europaeus* strains.

### 2.6. Characterization of the antibiotic resistance profile

Antibiotic resistance profiles were obtained based on the consensus obtained between the *in-silico* (i) and *in-vitro* (ii) analyses, and plotted in a presence-absence matrix using ComplexHeatmap 2.11.1:

i. Search for antibiotic resistance genes was performed using the web-based servers RGI 5.2.1 (McArthur, 2013) using the CARD database considering only strict and perfect matches (bitscore > 500), and ResFinder 4.1 (Bortolaia, 2020) with 80% of threshold and minimum length.
ii. Available bacterial strains (38 strains) were grown as described in 2.1. and antibiograms were carried out by the disc diffusion method on Müeller–Hinton agar (Oxoid) supplemented with 1% NaCl (MHA-1) with commercial discs (Oxoid, UK). The antibiotics tested were tetracycline (TE, 30 µg), oxytetracycline (OT, 30 µg), cephalexin (CN, 30 µg), ampicillin (AMP, 10 µg), amoxycillin (AML, 25 µg), erythromycin (E, 15 µg), enrofloxacin (ENR, 5 µg), flumequine (UB, 30 µg), cefoxitin (FOX, 30 µg), chloramphenicol (C, 30 µg), florfenicol (FFC, 30 µg), streptomycin (S, 10 µg), sulfonamide (SULDD, 25 µg) and gentamicin (CN, 10 µg). After incubation (24 h at 25°C), the zones of inhibition around the discs were measured and compared against recognized zone size ranges established by the manufacturer for each specific antimicrobial agent.

### 2.7. Search for secondary metabolite biosynthetic gene clusters

Biosynthetic gene clusters (BGC) were identified in each bacterial genome (39 genomes) using antiSMASH 7.0 web server (https://antismash.secondarymetabolites.org/) with default parameters and considering only strict findings. Genetic diversity within each BGC cluster among the 39 strain genomes was evaluated using BIG-SCAPE 1.1.5 (Navarro-Muñoz, 2020), including the determination of biosynthetic gene cluster families (GCF).

### 2.8. Characterization of anti-phage defense systems

All genome assemblies (39 assemblies) were individually analyzed using the following web-based servers: Defense-Finder (Tesson, 2022) and PADLOC v1.1.0 (v1.4.0) (Payne, 2022) with CRISPRDetect option (Biswas, 2016). A consensus between Defense-Finder and PADLOC outputs is shown with an absence-presence matrix and a PCA was performed with each strain profile as described in 2.2.

## 3. RESULTS

### 3.1. V. europaeus has an open pangenome dominated by the accessory genome

The features of *V. europaeus* assemblies used in this study are summarized in Supplementary Table 1. The *V. europaeus* pangenome was composed by a total of 9860 genes and it is considered as an open genome according to the Heaps value obtained (α=0.68) (Supplementary Fig. 1). Among these genes, 39% (3846 genes) were assigned to the core genome, whereas 61% (6014 genes) belonged to the accessory genome. Within the accessory genome, the 26%, 4% and 70% were assigned to the shell, soft, and cloud gene fractions respectively (Fig. 1A and 1B). In relation to the geographical origin, the main source of cloud genes were derived mainly from the French strains (CECT8427=666 genes; 07/115 T2==617 genes; 07/038 2T2=487 genes; 07/117 T1=468 genes; 07 108 T1=455 genes; CECT8426=321 genes; and 07/120 T1=316 genes), despite most of them were isolated from the same sampling time/location (Fig. 1B). A significant number of cloud genes were also associated to two Spanish strains (EX1=388 genes and PP2-843=320 genes) and in the American strain 071316F (542 genes) (Fig. 1B).

**Figure 1.**
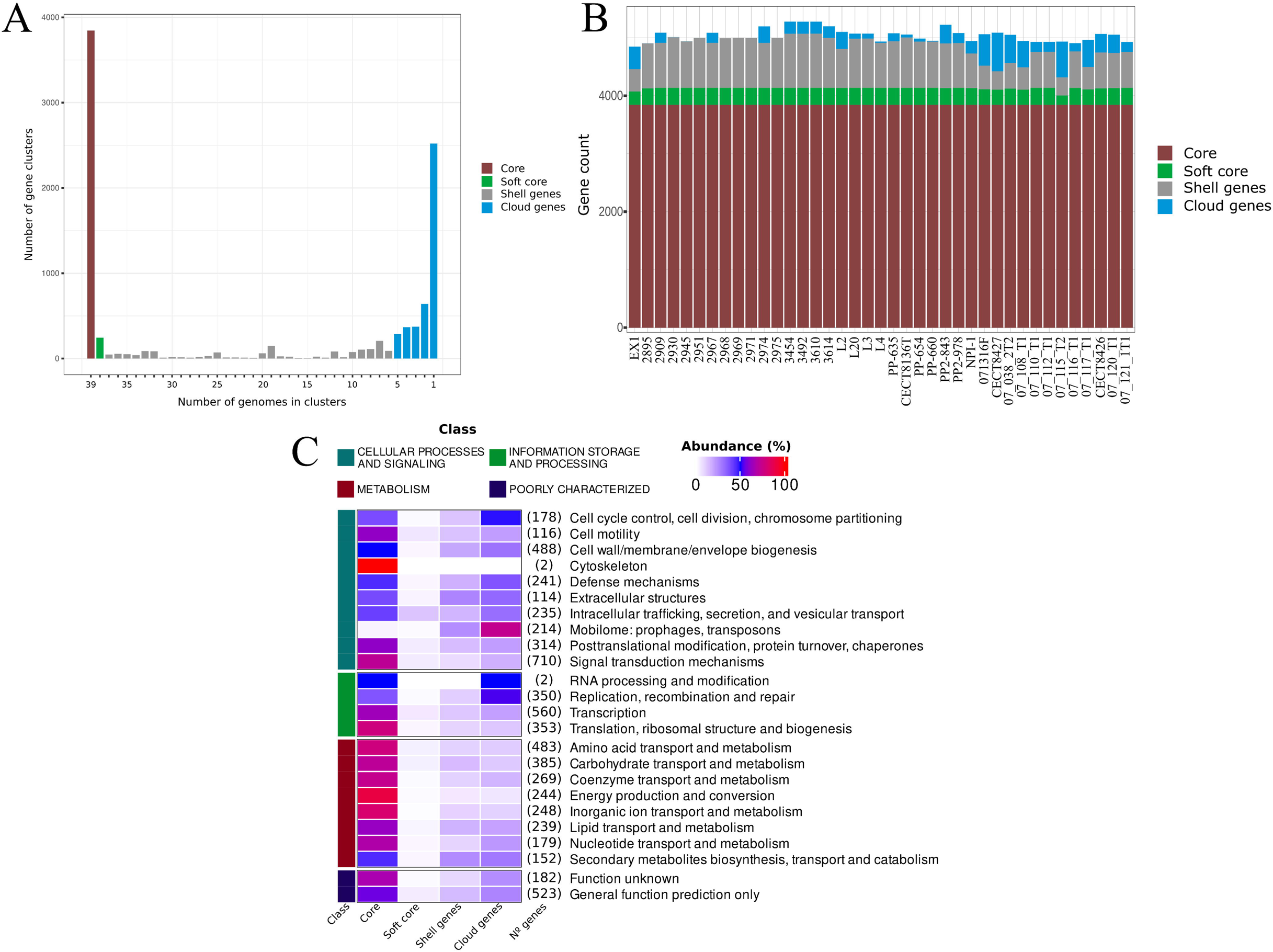
Main features of the *V. europaeus* pangenome. (A) Gene fraction (core, shell core, soft core and cloud) profiles. (B) Frequency distribution of gene clusters throughout the pangenome. (C) Percentage of genes related with COG general functions found in core, soft core, shell core, and cloud fractions of pangenome.

The 69% of the pangenome (6781 genes) was accurately assigned to specific COG functions, representing the 61%, 3%, 12% and 24% of the core, shell core, soft core, and cloud fractions, respectively (Fig. 1C). For the majority of the COG categories established, the genes identified were located into the core genome category (>50%) (Fig. 1C). Interestingly, the COG functions where accessory genes were predominant (>50%) always classified into the cloud category while they were almost absent in soft-core and shell genes. These more abundant categories in cloud genes were “Cell cycle control, cell division, chromosome partitioning”, “Defense mechanisms”, “Extracellular structures”, “Intracellular trafficking, secretion, and vesicular transport”, “Mobilome: prophages, transposons”, “Replication, recombination and repair” and “Secondary metabolites biosynthesis, transport and catabolism” (Fig. 1C).

Furthermore, differences in the number of RNA coding genes were identified comparing the six chromosome-level assembled genomes with the Illumina’s assemblies. Among the Illumina assemblies, tRNAs ranged from 88 (071316F) to 109 (2971 and 97/121 1T1) (Supplementary Table 2) with an average of 106 tRNAs/genome. However, the six highly contiguous chromosome-level assembled genomes encoded the higher number of tRNAs, ranging from 118 tRNAs (CECT8136 and NPI-1) to 122 tRNAs (CECT8427). Furthermore, rRNAs from Illumina assemblies displayed a lower copy number (1 copy for 16S rRNA, 1-4 copies for 5S rRNA and 1-2 for 23S rRNA) than the fully resolved genomes (9-10 copies for 16S rRNA, 10-11 copies for 5S rRNA and 9-10 copies for 23S rRNA) (Supplementary Table 3).

### 3.2. Nucleotide diversity and phylogenomic analyses based on the core-genome shed light on the evolutionary history and intraspecific diversity of V. europaeus

The phylogenomic tree based on the concatenated sequences of the 3846 core-genes showed a topology with three main branches containing a total of eight robust clusters (boostrap value=100; terminal or sub terminal branches distance > 0.006): branch I (cluster I), branch II (cluster II) and branch III (clusters III-VIII), being clusters I and II divided into sub clusters (boostrap value=100; terminal branches distances from 0.0003 to 0.0025) (Fig. 2). These phylogenetic relationships were supported by ANIb analysis, ANI values were higher than 0.979 in all pairwise comparisons, and around 0.995 for the strains belonging to the same cluster including between closely related sub-clusters (Supplementary Fig. 2). Interestingly, analyses of the phylogeny based on accessory genes showed the same 8 clusters although, as expected, the diversity was higher than the core genome phylogeny (Supplementary Fig. 3).

**Figure 2.**
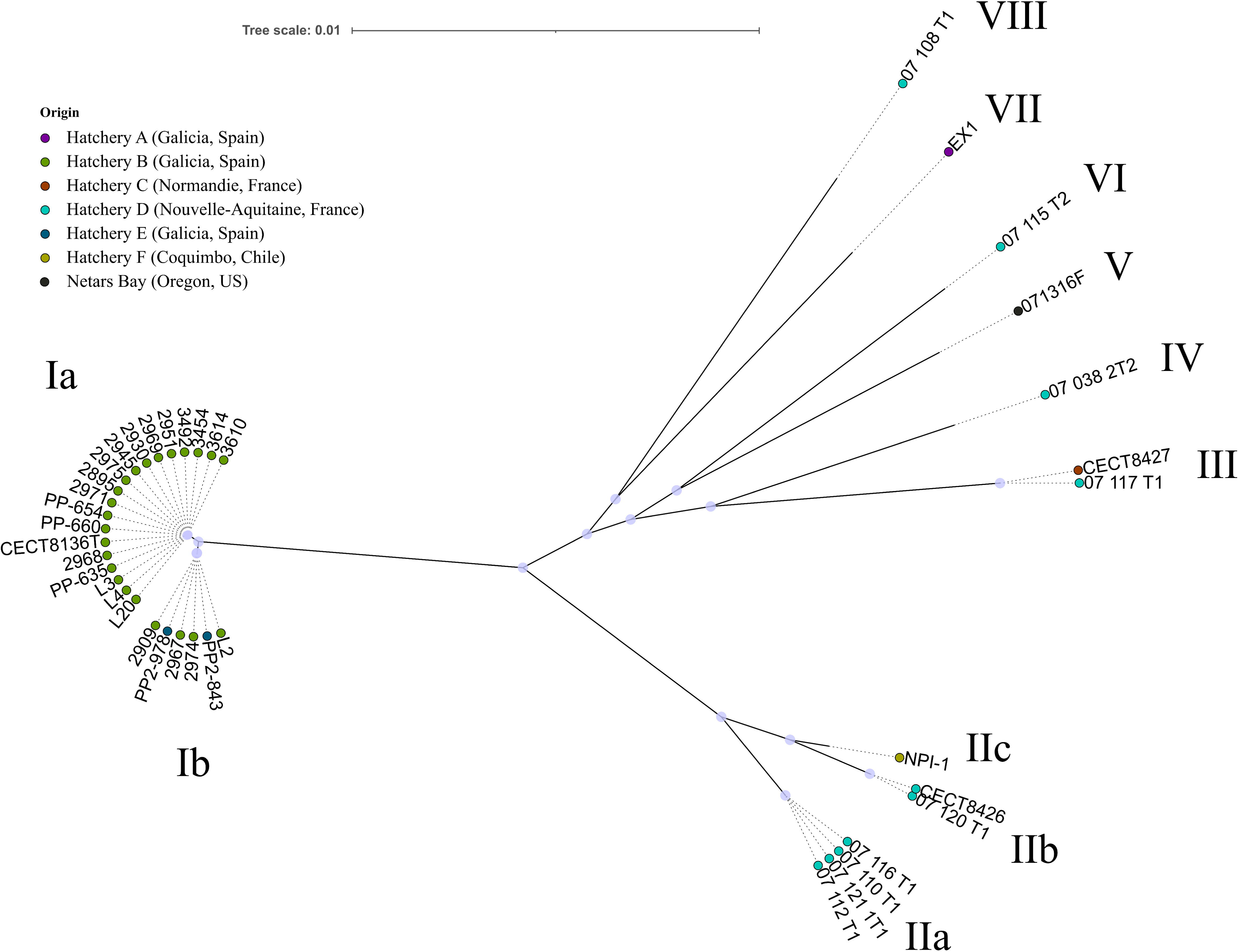
Phylogenenomic tree of *V. europaeus*. It was built by the concatenation of 3846 concatenated core-genes. Nodes with significant bootstrap values (n=100) are marked with a light purple dot. Clusters are indicated with roman numerals (I-VIII).

Cluster I was the largest group (25 strains), and it was exclusively constituted by the strains isolated from the Spanish hatcheries B (23 strains) and E (2 strains) (Table 1). Those strains were distributed into two sub-clusters: Ia (19 strains) and Ib (6 strains). The cluster comprises strains isolated in different years (2001, 2008, 2011 and 2018), from a broad host range and stages of the life cycle of bivalves such as eggs, larvae, spat -or juveniles- and adults (Fig. 2; Table 1). A complementary analysis to assess genetic differentiation and diversity within and between clusters was performed using SNPs detected in the coding regions of the core genome. Nucleotide diversity between sub-clusters Ia and Ib was low, including a maximum of 197 SNPs and 162 SNPs among the strains belonging to the sub-cluster Ia (CECT8136T vs. 2968) and sub-cluster Ib (2974 vs. L2) respectively, however more than 2300 SNPs were identified between both sub-clusters (Fig. 3A). Furthermore, analyses of the core-genome SNP dataset revealed the existence of many clonal Spanish strains (Table 2), isolated from: (i) from the same bivalve species (but from different samplings) into the sub-cluster Ia and sub-cluster Ib; (ii) broodstock to the offspring after spawning (vertical transmission); (iii) different hatcheries, hosts and/or dates. In contrast, there were SNP differences among strains isolated from the same sampling, despite they were assigned to the same sub-cluster, such as 89 SNPs between PP-635 and PP-660/PP-654 or 146 SNPs between PP-635 and CECT8136; or 118 SNPs between 2968 and 2895/2930/2945/2951/2969/2971/2975 (Fig. 3A).

**Figure 3.**
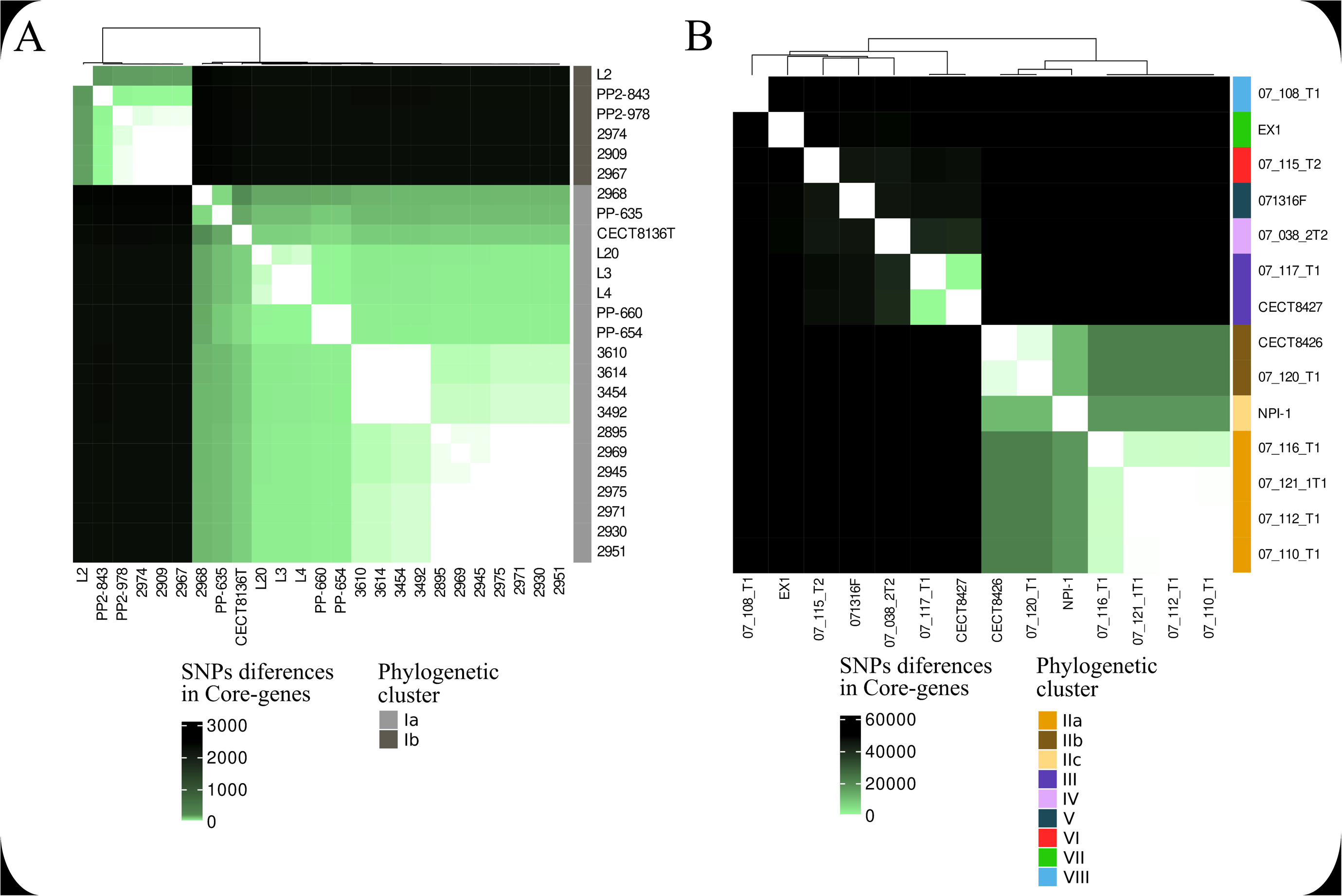
Heatmaps showing the genetic variability based on the SNP differences in the core-genes of the *V. europaeus* strains belonging: (A) Sub-clusters Ia and Ib; (B) Clusters from II to VIII. In both heatmaps, phylogenetic clusters and sub-clusters are shown in the right bar.

**Table 2.** Clonal strains identified from SNP analysis of the *V. europaeus* core-genome.

| Isolated from: | Location(s): | Host(s): | Cluster: | Clonal strains: | SNP difference: |
| --- | --- | --- | --- | --- | --- |
| Same bivalve species | Hatchery B (Spain) | *D. trunculus* (larvae and seawater) | Ia | 3454, 3492 | 0 |
|  | Hatchery B (Spain) | *R. decussatus* (larvae and seawater) | Ia | 2895, 2930, 2945, 2951, 2969, 2971, 2975 | 1-2 |
|  | Hatchery B (Spain) | *E. arcuatus* (larvae) | Ia | L3, L4 | 5 |
|  | Hatchery B (Spain) | *O. edulis* (larvae) | Ia | PP-654, PP-660 | 1 |
|  | Hatchery B (Spain) | *R. decussatus* (larvae) | Ib | 2909, 2974, 2967 | 1 |
|  | Hatchery E (Spain) | *R. philippinarum* (spat) | Ib | PP2-843, PP2-978 | 14 |
|  | Hatchery D (France) | *M. gigas* (spat) | IIa | 07/110 T1, 07/112 T1, 07/121 T1 | 2 |
|  | Hatchery D (France) | *M. gigas* (spat) | IIb | 07/120 T1, CECT8426 | 28 |
| Vertical transmission | Hatchery B (Spain) | *R. decussatus* broodstock to eggs | Ia | 3610, 3614 | 0 |
| Different hatcheries/hosts/dates | Hatchery E & B (Spain) | *R. philippinarum* vs. *R. decussatus* | Ib | PP2-978 vs. 2909/2974/2967 | 3 |
|  | Hatchery B (Spain) | *R. philippinarum* vs. *E. arcuatus* | Ia | L20 vs. L3/L4 | 5 |
|  | Hatchery B (Spain) | *D. trunculus* vs. *R. decussatus* | Ia | 3454/3492 vs. 3610/3614 | 1 |
|  | Hatchery B (Spain) | *R. decussatus* vs. *D. trunculus* | Ia | 2895/2930/2945/2951/2969/2971/2975 vs. 3454/3492 | 5 |
|  | Hatchery B (Spain) | *R. decussatus* (larvae) vs. *R. decussatus* (broodstock and eggs) | Ia | 2895/2930/2945/2951/2969/2971/2975 vs. 3610/3614 | 6 |
|  | Hatchery B (Spain) | *O. edulis* vs. *R. philippinarum* vs. *E. arcuatus* | Ia | PP-654/PP-660 vs. L20; PP-654/PP-660 vs. L3/L4 | 19 |

EX1 constituted the first known isolation of *V. europaeus* (year=1985) and the unique isolated from the Spanish Hatchery A (Table 1). This could support that EX1 was the only Spanish strain located out of cluster I and it constituted itself a well-differentiated group (cluster VII) in branch III, closer to the French strains than the Spanish strains (Fig. 2).

Conversely, most of the French strains (10/11 strains) were isolated from the same geographical origin and similar time (Table 1), however they exhibited much more genetic diversity than the Spanish strains (Fig. 2). Those ten French strains were distributed among six different clusters: branch II (cluster II=sub-clusters IIa, IIb and IIc) and branch III (clusters III, IV, VI and VIII) (Fig. 2). SNP analyses also revealed the existence of clonal French strains within the cluster IIa and IIb (Table 2). Cluster III (Fig. 2) was formed by two strains (07/117 T1 and CECT8427) from two different origins (Hatchery D and Hatchery C, located in Nouvelle-Aquitaine and Normandy respectively; Table 1) and hosts (*M. gigas* spat and the non-bivalve species *Haliotis tuberculata*; Table 1) identifying 182 SNPs among them (Fig. 3B). Other strains isolated from Hatchery D (07/138 2T2, 07/115 T2 and 07/108 T1) constituted by themselves independent clusters (clusters IV, VI and VIII) supported by differences of more than 50k SNPs with their contemporary strains belonging to sub-cluster IIa, IIb and cluster III (Fig. 3B).

Despite the Chilean strain NPI-1 was closely related to sub-cluster IIb, it constituted the sub-cluster IIc (Fig. 2) supported by a difference of 11.208 SNPs with its closest relatives (Fig. 3B).The American strain 071316F was the only one non-isolated from a hatchery environment and it also grouped in an independent cluster (cluster V), sharing a node with the French strain 07/115 T2 (Fig. 2).

### 3.3. The key virulence factors are encoded in the core-genome

According to the virulence challenges, all the *V. europaeus* strains tested (38 strains) were virulent against Manila clam juveniles (Fig. 4A). Survival rates were low in all cases, ranged from 0-7% in most cases (32/38 strains), 22-27% for 3/38 strains (CECT8427, 07/116 T1 and 2967) and 9-16% for 3/38 strains (07/121 1T1; 2968 and 2969) (Fig. 4A).

**Figure 4.**
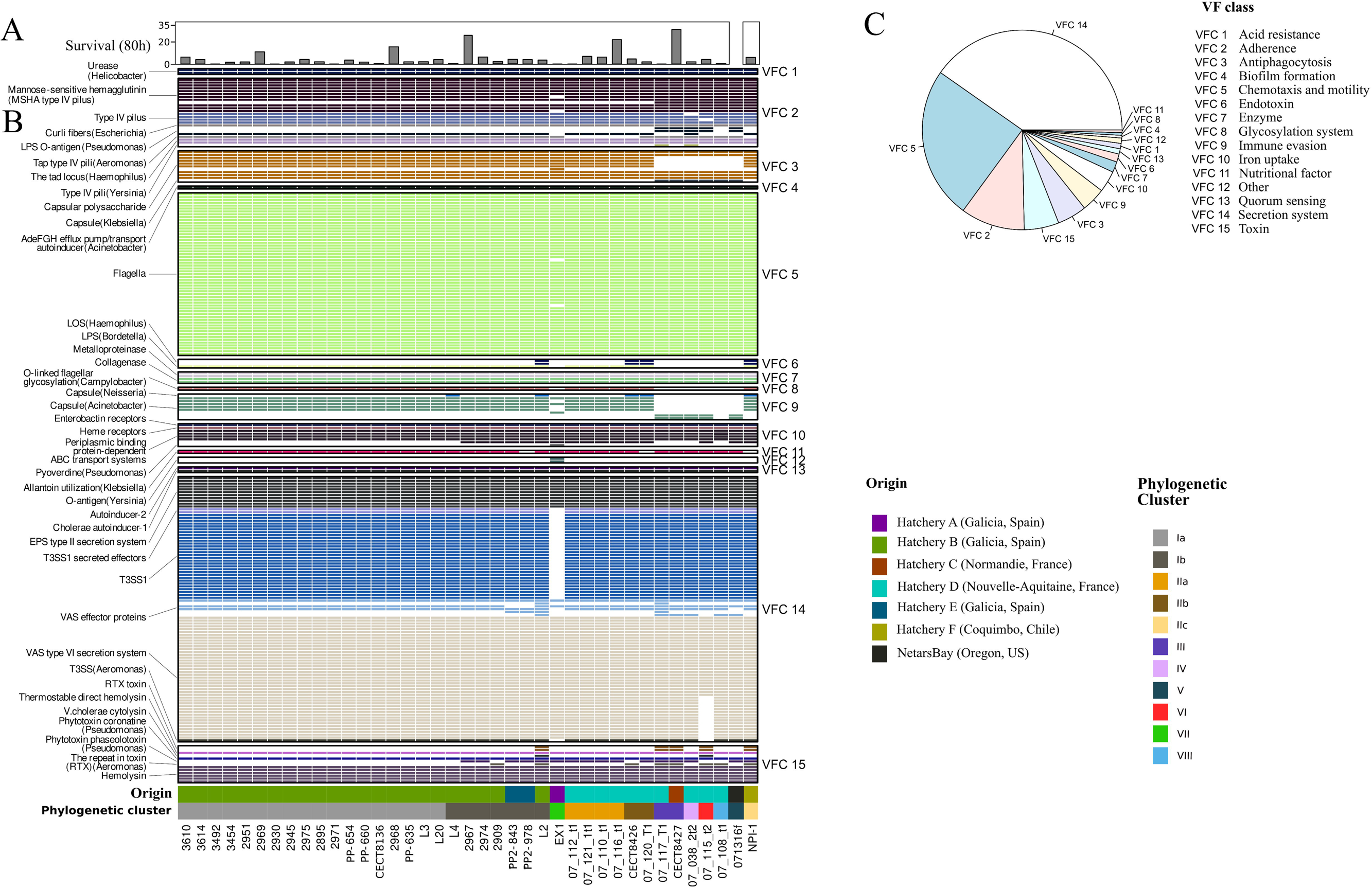
Virulence genes identified from the *V. europaeus* genomes: (A) Survival of Manila clam juveniles (%) after *V. europaeus* infection. (B) Presence/absence of virulence genes. (C) Relative abundance of the different Virulence Factor classes (VFCs) in the *V. europaeus* pangenome. Virulence factors are indicated on the left and VFCs on the right. Geographical origin of each *V. europaeus* strains and their phylogenetic clusters are indicated at the bottom.

A total of 231 genes (Fig. 4B; Supplementary Table 3) were identified as potential virulence factors from the *V. europaeus* pangenome and those were distributed in 15 Virulence Factor Classes (VFC1-15) (Fig. 4B and 4C). Among these genes, 59.3% (137 genes) were assigned to the core genome. Core virulence genes (Fig. 4B; Supplementary Table 3) were related to acid resistance (VFC 1), adherence (VFC 2), antiphagocytosis (VFC 3), biofilm formation (VFC 4), chemotaxis and motility (VFC 5), enzymes (VFC 7), iron uptake (VFC 10), quorum sensing (VFC 13), secretion systems (VFC 14) and toxins (VFC 15). In contrast, other VFC such as endotoxins (VFC 6), glycosylation system (VFC 8), immune evasion (VFC 9), nutritional factors (VFC 11) and others (VFC 12) did not contain any gene assigned to the core-genome.

It is important to remark that three classes, VFC 14, secretion systems; VFC 5, chemotaxis and motility; and VFC 2, adherence, comprised 75% of the virulence factors (Fig. 4B and 4C). Particularly, the pangenome included the whole gene machinery of five secretion systems (Fig. 4B; Supplementary Table 3): T2SS, T3SS and three different T6SS (T6SS1, T6SS2 and T6SS3). From those, the T2SS, T6SS1 and T6SS3 belonged to the core genome (Fig. 4B). T6SS2 and T3SS were assigned to the accessory genome because despite they were encoded by most of *V. europaeus* strains and only the French strain 07/115 T2 was defective for T6SS2 and the Spanish strain EX1 for T3SS (Fig. 4B). Despite both T6SS1 and T6SS2 showed a similar synteny and belonged to the T6SSi5 subtype, they presented significant differences among gene homologues (Supplementary Fig. 4). T6SS3 is different from 1 and 2, it was classified into the T6SSi1 subtype (Supplementary Fig. 4).

The PCA analysis (Fig. 5A) explained the 85.16% of the genetic diversity of the non-core virulence factors. Thereby, genetic variability based on the non-core virulence factors was low and they showed differences only in the presence/absence of few virulence genes among closest relatives (Supplementary Table 4). The co-existence of different virulence profiles in the same environment could be evaluated in the Hatchery B, in this case the strains obtained in this facility isolated from different cultures along the time-series presented different profiles (2001-2018) (Fig. 5A). Five different virulence factors profiles were identified in this Hatchery (VFP1-5), although they were very similar differing mostly in single genes (Fig. 5A). For instance, VFP 1 was always detected along the time-series, being the unique profile in Mar/2001, May/2012, Jun/2012 and Jul/2012 and those strains shared 70 non-core virulence factor genes. VFP 1 co-existed with VPF 3 and VFP 4 in May/2011 and with VFP 2 and VFP 5 in Mar/2018 (Fig. 6A). Interestingly, the strain L2 represented the VFP4 and it harbored the highest number of virulence genes (82 genes) among the *V. europaeu*s strains (Fig. 6A).

**Figure 5.**
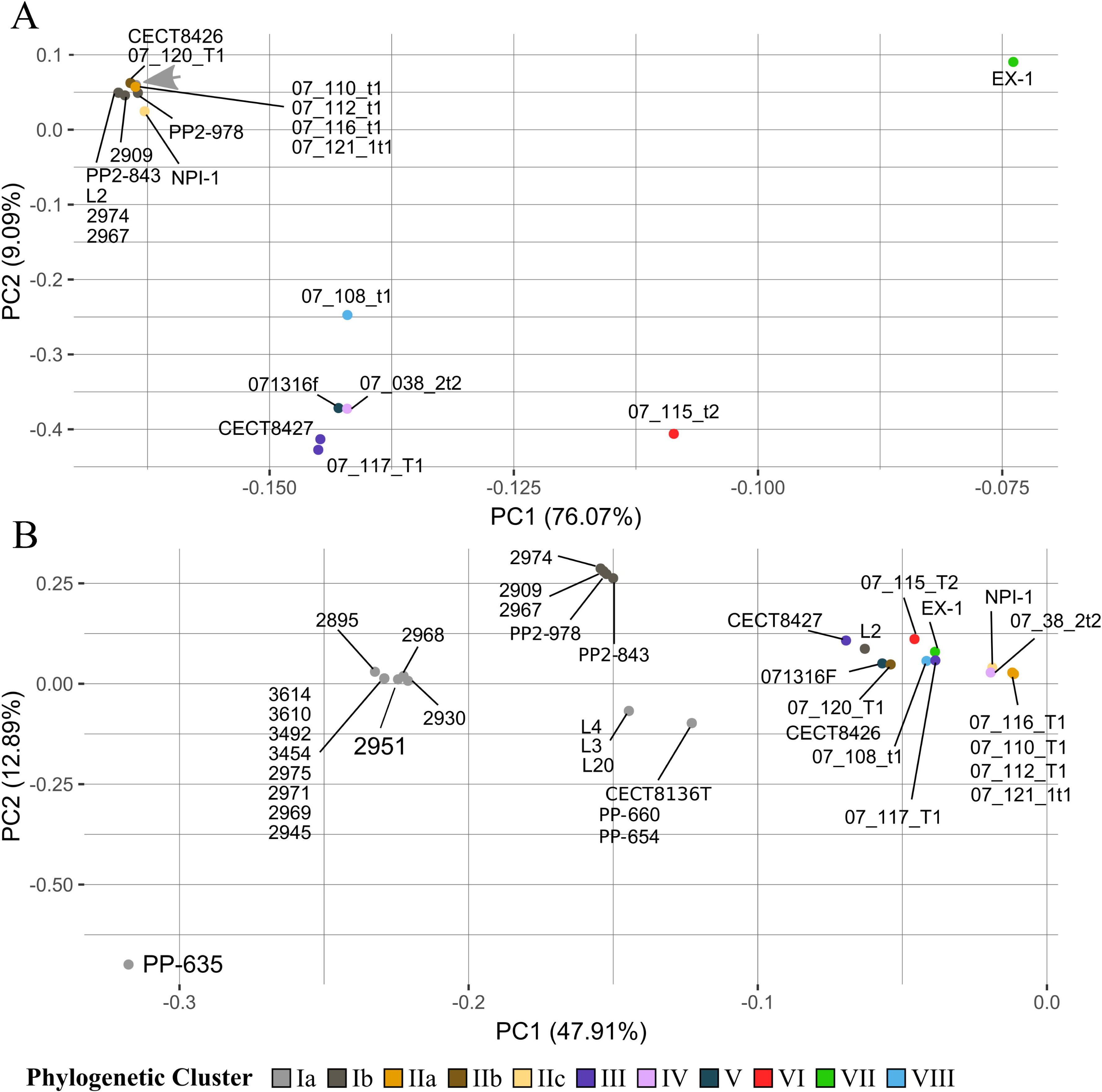
Genetic variability of *V. europaeus* strains based on the virulence factors (A) and anti-phage defense systems (B) encoded in the accessory genome. Phylogenetic clusters based on the core-genome were shown in different colors in PCA. Grey arrow in indicates the position in the PCA of the strains: PP-660, PP-635, CECT8136, PP-654, 2895, 2945, 2968, 2930, 2951, 2969, 2671, 2975, 3610, 3492, 3454, 3614, L3, L4 and L20.

**Figure 6.**
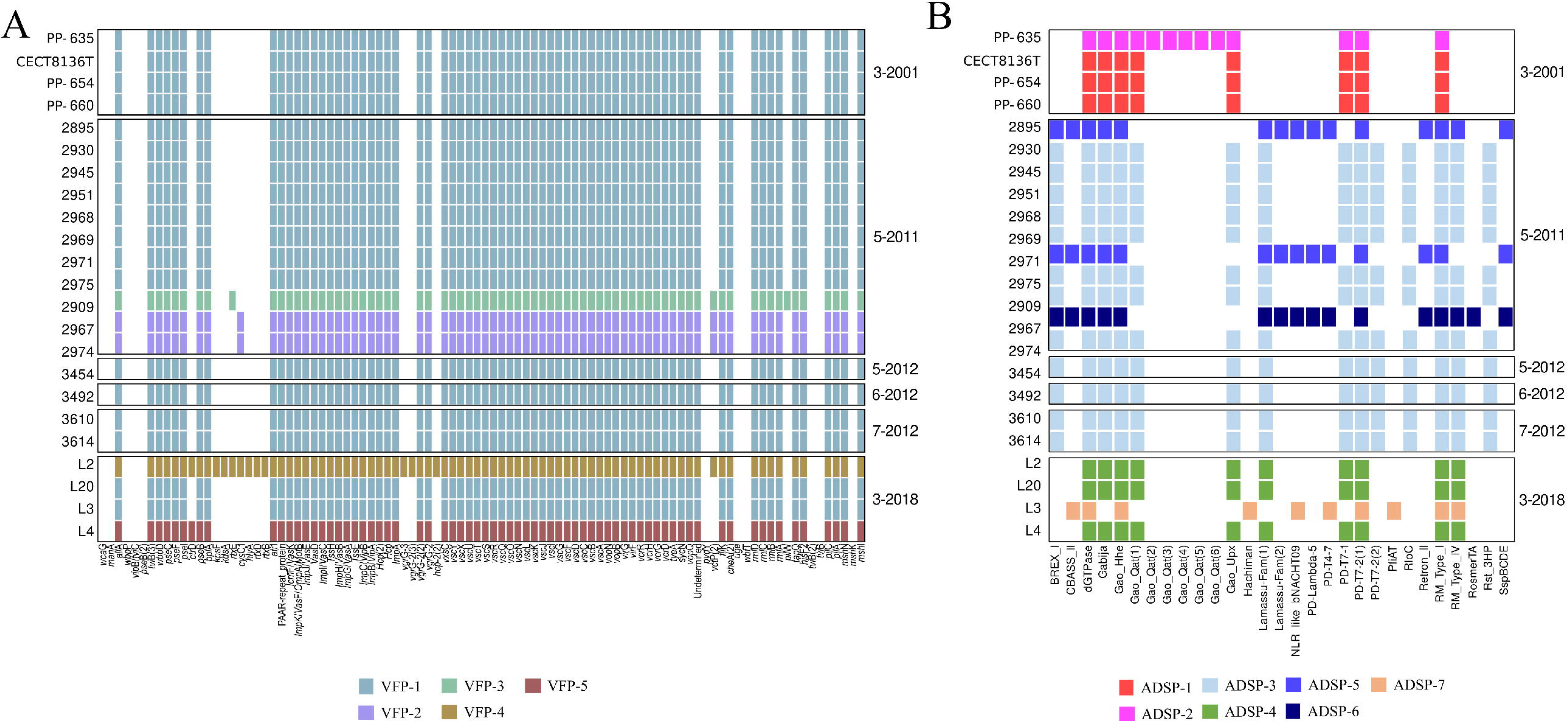
Co-existence of different virulence factors profiles (A) and anti-phage defense systems (A) in the Hatchery B (Spain) along the time series (2001-2018). Presence/absence of accessory virulence-related genes were displayed in rows and the different profiles (VFP1-5 or ADSP 1-7) indicated by colors. (A) Virulence factor profiles (VFP 1-5), VPF 2 and VPF 3 were very similar to VFP1 however they differed in the presence of VAS effector protein coding genes *vgrG-2*, the allantoin utilization gene *allA* and the presence/absence of some toxins such as CysC1 (present only VPF 3) or RtxE (present only in VPF 2). VFP5 was identical to VFP1 but it encoded an additional non-core immune evasion gene *ctrD*. VFP4 included 12 additional non-core virulence genes in comparison with VFP1 such as those encoding an additional VAS effector proteins of the T6SS encoded in the accessory genome (*hcp-2_2*, *vgrG-2*, *vgrG-2_3* and *vgrG-3*), toxins (*rtxB*, *rtxD*, *rtxE*, *hlyA* and *cysC1*), endotoxins (*kdsA* and *kpsF*), immune evasion (*ctrD*) and allantoin utilization (*allA*). (B) Anti-phage Defense Systems Profiles (ADSPs 1-7) included the following systems: ADSP 1 (dGTPase, Gabija, Gao Hhe, Gao Qat, Gao Upx, PD T7 1, PD T2 7 and RM type I); ADSP2 was similar to ADSP1 but ADSP2 harbored five additional copies of the Gao Qat system; ADSP 3 was similar to ADSP 1 encoding additional defense systems such as BREX I, Lamassu Fam, Rloc, RM type IV and Rst_3HP; ADSP 3 was similar to ADSP 1 encoding also Lamassu Fam and RM type IV; ADSP 5 and ADSP 6 shared 14 defense systems BREX I, CBASS II, dGTPase, Gabija, Gao Hhe, Lamassu Fam (two copies), NLR like bNACHT09, PD lambda 5, PD T7 2, Retron II, RM type I, RM type IV and SspBCDE but ADSP5 (2909 and 2967) also encoded PD T4 7. ADSP6 (2974) encoded additional systems such as PD T4 7 and RosmerTA; ADPS7 (L2) showed a very different profile than its contemporary strains L3, L4 and L20, encoding a total of eight defense systems (CBASS II, dGTPase, Gao Hhe, Hachiman, NLR like bNACHT09, PD T4 7, PfiAT, and RM type I) and it was the only one encoding the PfiAT system.

French strains showed a higher variability based on non-core virulence genes than the Spanish strains but lower than the observed in the core-genome phylogeny. For instance, strains belonging to sub-cluster IIa (07/112 T1, 07/121 1T1, 07/110 T1, and 07/116 T1) were close to Spanish strains (sub-cluster Ia and Ib) (Fig. 5A). Interestingly, strains 07/115 T2 (cluster VI) and EX-1 (cluster VII) showed the most different virulence profiles (Fig. 5A) because they encoded a lower number of non-core virulence genes (07/115 T2= 57 genes; EX1= 38) than their closest relatives since they were defective for T6SS2 and T3SS, respectively.

### 3.4. Antimicrobials are (still) effective to fight V. europaeus

A consensus between the *in silico* and phenotypic results was established to determine the antibiotic resistance profile. *V. europaeus* was sensitive to most of the antimicrobials evaluated in this study. All *V. europaeus* strains available in our lab collection (38 strains) were resistant to erythromycin (E, 15 μg) according to the disc diffusion method, which was confirmed with the presence of the Antibiotic Resistance Gene (ARG) *crp* (Supplementary Fig. 5). The strains 3454, 3492, 3610 and 3614 were also resistant to streptomycin (S, 10 μg) and sulfonamide (SULDD, 25 μg) bestowed by the presence of the ARGs *aph(3’’)-Ib* and *aph(6)-Id* genes and *sul2* gene, respectively, in their genomes (Supplementary Fig. 5).

Some discrepancies were found between *in silico* and phenotypic results, and thus *in silico* data was curated with the results obtained by the disc-diffusion method. For example, despite no cephalosporin resistance gene was found in CARD and ResFinder databases, all *V. europaeus* strains tested were experimentally resistant to cephalexin on MHA-1 plates. On the other hand, some resistance genes found by *in silico* analyses rendered a negative phenotypic result: (i) *floR* gene was identified in the genomes of the strains 3454, 3492, 3610 and 3614 by ResFinder (identities ranged from 98.19%-98.27%), however, they were sensitive to florfenicol (FFC, 30 μg); (ii) tetracycline resistance genes were also found in all strains from both databases (gene identity >85%), however they were sensitive to tetracycline (TE, 30 μg).

### 3.5. V. europaeus produces an important number of secondary metabolites mostly encoded in the accessory genome

*V. europaeus* genomes encoded a total of 254 Biosynthetic Gene Clusters (BGCs). Most of the strains encoded 6-7 BGCs, ranging from the 5 BGCs encoded by the French strain 07/115 T2 (cluster VI) to the 8 BGCs of the American strain 071316F (cluster V) (Fig. 7). Study of the intra-BGC diversity allowed its classification in four major classes encoding the following BCGs (%): PKS-NRP hybrids (14.57%), RiPPs (15.75%), NRPS (15.75%), and Others (53.94%). Within those classes, 12 biosynthetic Gene Clusters Families (GCF) could be defined (Fig. 7): GCF1 and GCF2 presumably produced PKS-NRP hybrids; GCF3 and GCF4 post-translationally modified peptides (RiPPs); GCF5 and GCF6, non-ribosomal peptide synthetases (NRPS); GCF7 and GCF8, ectoine; GCF9, arylpolyene-NRPS hybrids; GCF10, butyrolactone; GCF11, betalactone; GCF12, arylpolyene.

**Figure 7.**
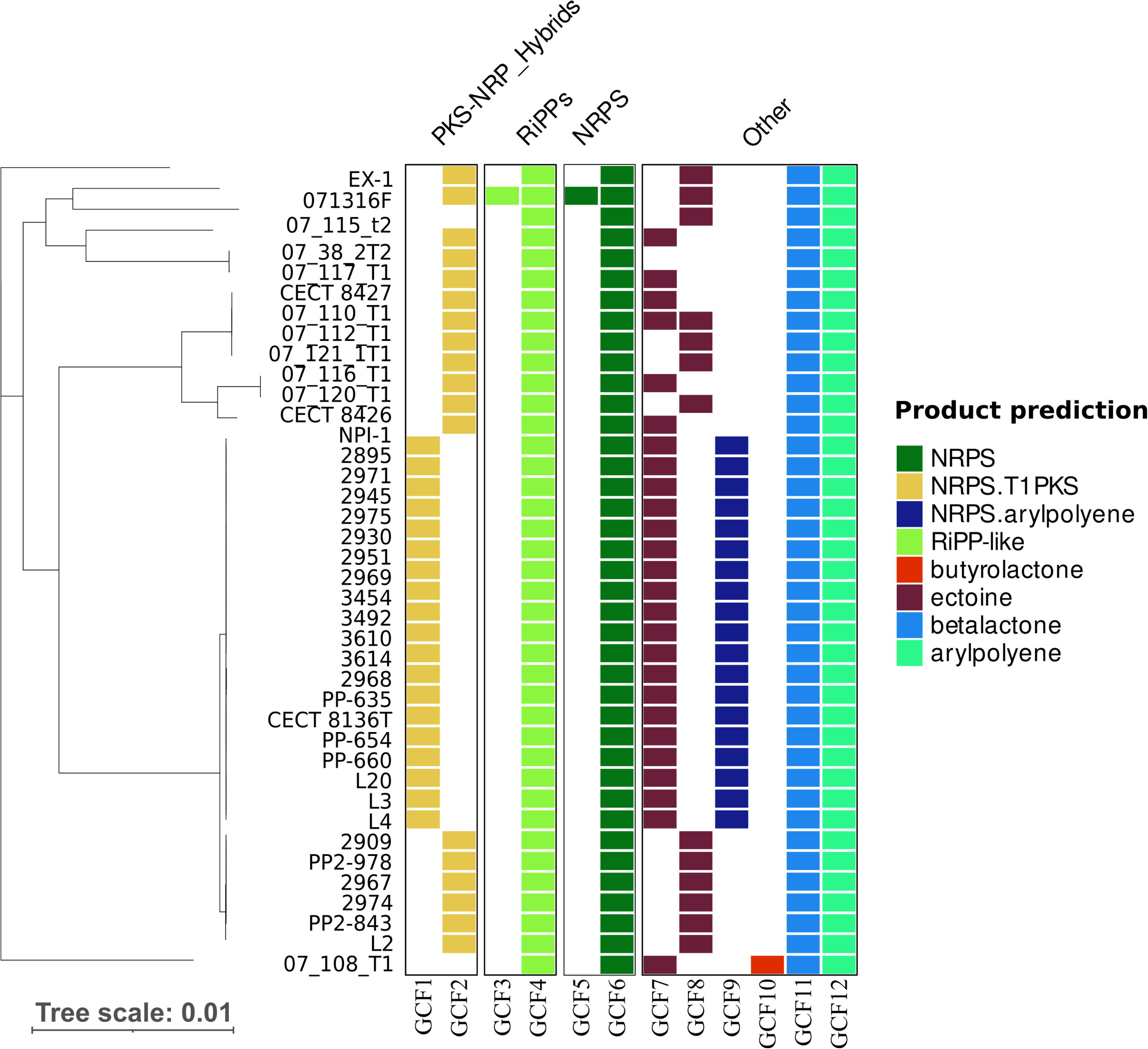
Heatmap showing the biosynthetic gene cluster families (GCFs) encoded by each V. europaeus strain. Products from each GCF are indicated by different colours. Core-genome phylogeny was plotted on the left

GCF4, GCF6, GCF11 and GCF12 contained BGCs encoded by all *V. europaeus* strains and, thus, they were assigned to the core-genome (Fig. 7). GCF1, GCF2, GCF7, GCF8, and GCF9 are conformed by sequences of just a subset of strains (Fig. 7): GCF1 and GCF9, encoded only by the Spanish strains belonging to sub-cluster Ia; GCF2, encoded by Spanish, French, Chilean and American strains (clusters Ib, IIa-c, III, IV, V, and VII); GCF7, encoded by Spanish, French and Chilean strains (clusters Ia, IIa-c, III, VIII, and IV); GCF8, encoded by Spanish, French and American strains (clusters Ib, IIa-b, V, VI, VII). Three GCFs were encoded by a single genome: GCF3 and GCF5 were only encoded by the American strain 071316f (US; phylogenetic cluster V), whereas GCF10 was only detected in the French strain 07/108 T1. According to antiSMASH results, GCF5 presents a 100% similarity to the siderophore amphibactin B biosynthetic gene cluster of *V. neptunius*, and GCF12 presents an 85-90% similarity to arylpolyene Vf BGC of *Aliivibrio fischeri* ES114.

### 3.6. Variability of V. europaeus is explained by the presence of anti-phage defense systems in the accessory genome

*V. europaeus* genomes were enriched in anti-phage defense systems: a total of 49 systems were identified among the studied genomes (Fig. 8). It is important to remark that anti-phage defense systems were mostly encoded by the accessory genome and only the dGTPase system was identified into the core. Interestingly, Gao Hhe system was absent only in the French strain 07/115 T2 (Fig. 8).

**Figure 8.**
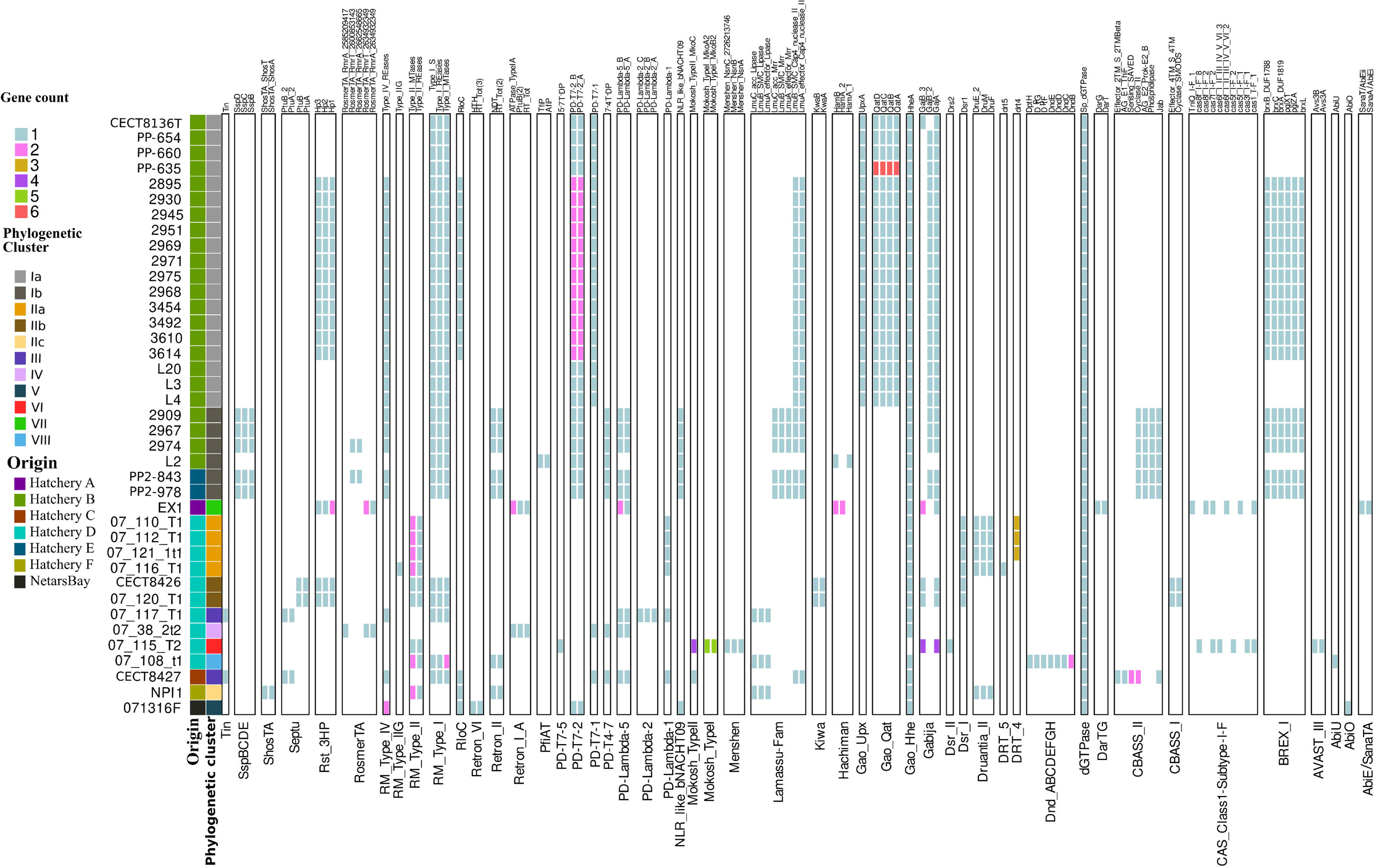
Anti-phage defense systems encoded by *V. europaeus* strains. Components are coloured by the number of copies and grouped by a specific defense system. Geographical origin of the *V. europaeus* strains, the phylogenetic cluster based on the core genomes and the anti-phage defense system profiles (ADSPs) are indicated on the left.

PCA analyses (Fig. 5B) revealed indeed the genetic variability among *V. europaeus* strains is mainly determined by the anti-phage defense systems in comparison with other accessory genes such as virulence genes, ARG or BGCs. For instance, different Anti-phage Defense Systems Profiles (ADSPs) coexisted to effectively protect the *V. europaeus* population from the phage predation in the Hatchery B and ADSP variability along the time series was higher than the observed for the virulence profiles (5 different VFP types vs 7 different ADSP types) (Fig. 6A and 6B). However, it is important to remark that anti-defense systems are formed by several genes in most cases. In Hatchery B, ADSP 1 and ADSP 2 profiles co-existed in Mar 2001 (Fig. 6B). Defense systems encoded by ADSP 1 were common in other new profiles such as ADSP 3 and ADSP4 (Fig. 6B). ADSP 3 was unique in strains isolated 2012 and it coexisted with ADSP 5 and 6 in May 2011 (Fig. 6B). Interestingly, ADSP5 (strains 2909 and 2967) was the most protected phenotype since it included the highest number of anti-phage defense systems with a total of 16 full systems. Besides, ADPS7 (strain L2) coexisted with ADSP 4 and showed a very different profile than its contemporary strains L3, L4 and L20 (Fig. 6B).

French strains isolated from hatchery D also showed high variability, supporting the coexistence of different profiles in the same hatchery (Fig. 5B). Within the sub-cluster IIa, three strains 07/110 T1, 07/112 T1 and 07/121 1T1 encoded seven systems such as dGTPase, and three copies of Drt 4, Druantia II, Dsr I, Gao Hhe, PD lambda I and RM type II. The remaining strain belonging to this sub-cluster (strain 07/116 T1) showed a similar profile, however, it harbored two additional systems being the only *V. europaeus* strains encoding the DRT 5 and RM type IIG systems (Fig. 8). Strains belonging to sub-cluster IIb (strains CECT8426 and 07/120 T1) encode 10 systems (CBASS I, dGTPase, Dsr I, Gabija, Gao Hhe, Kiwa, RM type I, RM type II, Rst 3HP and Septu). The remaining strains isolated from Hatchery D (Fig. 8) such as 07/117 T1 (cluster III), 07/038 2T2 (cluster IV), 07/115 T2 (cluster VI) and 07/108 T1 (cluster VIII) encoded exclusive defense systems, for example: (i) PD-Lambda-2 system only encoded by 07/117 T1; (ii) Retron IA system only encoded by 07/038 2T2, which it was the strain with the lowest number of defense systems with only six full systems; (iii) AVAST III, Dsr II, Menshen and Mokosh_Type I only encoded by 07/115 T2; (iv) AbiU and Dnd ABCDEFGH systems only encoded by 07/108 T1.

The remaining *V. europaeus* strains encoded exclusively specific defense systems. For instance, the Spanish strain EX1 (cluster VII), isolated from Hatchery A, was the only one encoding the DarTG and SanaTA systems (Fig. 8). Whereas, the only French strain isolated from a different hatchery (strains CECT8427; Hatchery C, Normandy; Fig. 8) encoded 11 defense system (CBASS II, dGTPase, Gao Hhe, Lamassu Fam, PD-lambda 1, PD-lambda 5, PD-T7 1, RloC, RM_Type_I, RM Type IV and Septu) and it was the only one with two copies of the genes of CBASS (Fig. 8). The American strains NPI-1 (Chile; cluster IIc) and 071316F (US; cluster V) encoded for a total of 7 and 9 defense systems respectively. Interestingly, some defense systems were only harbored by those strains such as AbiO and Retron VI systems only encoded by the strain 071316F or ShosTA system by the Chilean strain NPI-1.

## 4. DISCUSSION

The pangenome obtained for different species of the genus *Vibrio* showed a broad size range (Dias, 2018; Hansen, 2020; Guardiola-Avila, 2021; Zheng, 2022; Du, 2022), reflecting a high genomic diversity within the genus, which matches with its well-known ecological diversity (Nathamuni, 2019; Sampaio, 2022). The pangenome of *V. europaeus* (9860 genes) was comparable to other *Vibrio* pangenomes such as the multi host pathogens *V. anguillarum* (9537 genes) and *V. tapetis* (11213 genes) (Dias, 2018; Hansen, 2020). Regarding the percentage of core genes in the pangenome, it is strongly related to evolutionary history and lifestyle of a bacterial species. For instance, free-living species with high degree of dispersion have a small proportion of core genomes (McInerney, 2017). In our study, *V. europaeus* exhibited higher core genome (39.00%) than other *Vibrio* pathogens with a broad host-range, such as *V. anguillarum* (28.24%), *V. tapetis* (29.90%) or *V. fluvialis* (15.36%) (Dias, 2018; Hansen, 2020; Zheng, 2022). This can be justified due to the bias of the *V. europaeus* collection because most of the strains were isolated: (i) from marine mollusks but not from other Phylum; (ii) associated to massive mortalities but not from healthy animals; and (iii) from a reduced number of locations. Interestingly, a difference on the abundance of structural RNAs (tRNA and rRNA) were found between the six long-read sequencing assemblies and those sequenced with short-read Illumina technology. The six chromosome-level assembled genomes showed the highest number of RNAs, probably due to the limitations of short-reads to identify different genes of the same family, especially if they are tandemly arranged (Miyamoto, 2014).

Analysis of the core genome allowed us to evaluate the evolutionary history and intraspecific diversity of *V. europaeus*. Specifically, the Spanish strains belonging to phylogenetic sub-clusters Ia and Ib displayed a lower genetic variability than the French strains. As observed previously by Campbell, (2024) for the radiation event of the clonal type of *V. parahaemolyticus*, those Spanish populations spreaded from a monophyletic radiation event that it was recently dispersed artificially or naturally throughout the hatcheries under study (Hatchery B and E). In contrast, French strains exhibited a higher variability than the Spanish strains, even though the geographical and temporal distribution of its isolation were more restricted in the time and geographical location (Hatchery D and samplings performed in August 2007). In the case of Chilean strain NPI-1, its closeness to the French strains of sub-cluster IIb strongly suggest an intercontinental transference of *V. europaeus*, probably mediated by the anthropic movement (mollusks spats, broodstock, phytoplankton…). Besides for strain EX1 reflects historical practices in hatchery management, during that period broodstock were mostly imported from France facilitating the introduction of genetically diverse *V. europaeus* lineages (personal communication Juan L. Barja). This explains the distinct placement of strain EX1, clustering closer to French isolates rather than Spanish strains. This anthropic movement events would also support the finding of clonal strains in different locations in Spain and France.

Comparative analyses revealed that 60% of the virulence genes of the *V. europaeus* pangenome belonged to the core genome. It is important to remark that all *V. europaeus* strains were demonstrated to highly virulent towards Manila clam juveniles, in consequence, this led to the determination that the key virulence factors are necessarily encoded by the core genome. In the case of *Vibrio* pathogenic for bivalves, pathogenesis is multifactorial and it depends on the combination of different virulence factors, such as successful chemotaxis, adherence and a first colonization of bivalve tissues; survival to bivalve immune system and proliferation in hemolymph; colonization of the connective tissue, bacterial proliferation and nutrient acquisition, which finally causes tissular disruption and host death (Parizadeh, 2018; Destoumieux-Garzón, 2020). It is important to note that virulence factors of *V. europaeus* were strictly analyzed here by comparative genomics. However, genetic manipulation will be essential to demonstrate the functional role of those genes in virulence. Due to their presence in all *V. europaeus* strains, key pathogenic effectors seem to be driven by the core-secretion systems T2SS and both T6SS (T6SS1 and T6SS3). Virulence-related functions of T2SS includes the participation in attachment, biofilm formation and colonization, as well as the secretion to the extracellular medium proteins capable to produce tissue disruption including proteases, pectinases, phospholipases, lipases, and toxins (Sandkvist 2001; Cianciotto and White 2017). The T6SS plays a major role in interbacterial competition and in bacterial interactions with eukaryotic cells (Smith, 2020; Feria and Valvano 2020). Interestingly, 97% of the *V. europaeus* strains encoded three different T6SSs (T6SS1-3), which is an unusual phenomenon. T6SS1 and T6SS3, belonging to the core-genome and encoded by the Chromosome 1, appeared to be widespread among all the stains. In contrast, T6SS2 was assigned to the accessory genome and, thus, it was probably acquired later in a radiation by a horizontal gene transfer event or to a specific gene loss into that phylogenetic group (Murray et al 2020; Jana,, 2022; Morgado and Vicente, 2022). T6SS2 and T6SS3 are similar to the previously reported T6SS3-like of *V. proteolyticus* and *V. parahaemolyticus* involved on delivering anti-eukaryotic effector proteins into mammalian phagocytic cells (Jana,, 2022; Cohen,, 2023). T6SS1 is similar to the T6SS1-like of *V. coralliilyticus* which mediates antibacterial activities (Mass,, 2024ab). Our findings revealed a diverse repertoire of T6SSs in the *V. europaeus* pan-genome which can play a major role in the interactions of this species with other cells. On the other hand, the T3SS, a recognized virulence factor of Gram-negative bacteria capable of injecting effectors of variable functions into host cells (Cornelis and Van Gijsegem 2000; Park, 2004; Osorio 2018), is present in all the *V. europaeus* strains except for the oldest isolate (strain EX1), suggesting a recent horizontal acquisition of that operon (Brown and Finlay 2011). The T3SS identified in *V. europaeus* corresponded to the T3SS1 described in *V. parahaemolyticus*, this has been reported to be mainly related to biofilm formation, motility and cytotoxicity (Hiyoshi, 2010; Calder, 2014). Due the remarkable absence of T3SS in the pathogenic strain *V. europaeus* EX1, its role in the pathogenicity of the species deserves further investigation. In relation to other virulence factors, *vemA* and *prtV* genes (encoding M4 metalloproteinases), *colA* and *colP* (encoding for collagenases) and the seven hemolysin toxins were encoded in the *V. europaeus* core-genome. Metalloproteinases VemA and PrtV and their homologues were related with proteolysis and host’s colonization in a broad range of *Vibrio* taxa, such as *V. cholerae*, *V. anguillarum*, *V. aestuarianus*, *V. coralliilyticus*, *V. neptunius*, and *V. splendidus* (Vaitkevicius, 2006; Binesse, 2008; Hasegawa, 2008; Varina, 2008; Mo, 2010; Galvis, 2021). In a previous study, Martínez, (2022) determined that a mutant of *V. europaeus* CECT8136 defective in the *vemA* gene resulted in a slightly slow pathogenic process in larvae and juveniles of Manila clam and displayed a chemotaxis ability favored by VemA to colonize the body mucus of clams and to form biofilm.

All *V. europaeus* strains were resistant to cephalexin and erythromycin and they were sensitive to most of the antibiotics tested. Previous studies demonstrated the high prevalence of macrolides and cephalosporins antibiotic resistance genes among *Vibrio* species, including the aquaculture pathogen *V. parahaemolyticus* (Albini, 2022). In relation to the ARGs, the genetic basis of resistance to cephalosporins has not been identified in *V. europaeus* and could be due to specific mutations (Palace, 2020). It is important to note that four *V. europaeus* strains isolated from the Spanish Hatchery B were also resistant to two additional antimicrobials. Hatcheries and bivalves are optimal environments to promote the bacterial acquisition of ARGs by HGT due (i) to the extended use of antimicrobials to prevent or control bacterial diseases; and (ii) the bioaccumulation of antimicrobials in bivalve tissues (Dubert, 2017b; Baralla, 2021; Kijewska, 2023). This highlights the risk of using antibiotics in shellfish aquaculture despite most of the antibiotics tested could be useful to fight *V. europaeus* (Dubert et al 2017b; Baralla, 2021; Kijewska, 2023). From an environmental perspective, secondary metabolites are connected to the capacity of the bacteria to occupy its ecological niche, facilitate access to a specific nutrient, competing for resources or establishing relationships with other microorganisms (Giubergia, 2016; Modolon, 2020). Most of the secondary metabolites produced by *V. europaeus* (NRPS, RiPP-like, betalactone, arylpolyene, and ectoine) were found in other marine vibrios (Burks, 2017; Alex, 2021). A highly conserved amphibactin system was found in *V. europaeus* and this metabolite related to the capacity to acquire iron is widespread in commensal and pathogenic *Vibrio* associated to bivalve hemolymph (Galvis, 2020). The role of ectoine and arylpolyene could be related to bacterial fitness in osmotic and oxidative stress (Cimermancic, 2014; Czech, 2018). The ecological role of the RiPP (GCF4), the arylpolyene (GCF12) and the PKS-NRP hybrid families (GCF1 and GCF2) must be elucidated due to its high prevalence in the *V. europaeus* strains.

Our results demonstrated that intra-specific variability is mostly due to the presence of the most anti-phage defense systems in the accessory genome. This sheds light on the vast -and diverse-repertoire of anti-phage defense systems encoded by *V. europaeus*. According to Hussain, (2021), anti-phage protection is cumulative, and those defense systems constitute a large fraction of the accessory genome, accounting even for more than 90% of the accessory genome among closest relatives. This means the phage defense elements can evolve and transfer from cell to cell without interfering with metabolic or physiological processes encoded by the core genome, maintaining the core genome over the long term even in the face of phage predation (Hussain, 2021). The association of anti-phage system genes to the accessory genome reflects the host’s adaptation in the arms-race phage-bacteria. This idea is supported by the presence of multiple defense profiles in a close temporal and physical space, especially in clonal strains or strains belonging to the same phylogenetic cluster (Bernheim and Sorek 2020; Rocha and Bikard 2022; Botelho, 2023ab). The repertoire of defense systems is known as defensome and its understanding is important when considering the therapeutic use of phages in aquaculture. In this sense, phage therapy could not be effective against *V. europaeus* due to the important and diverse defense arsenal encoded by its accessory genome. Some authors have demonstrated that the rapid acquisition of bacterial resistance against phages offers a parallel to the spread of antibiotic resistance on plasmids in bacteria (Hussain, 2021). Thus, the routine administration of a phage cocktail in shellfish aquaculture to prevent bacterial diseases could not be the optimal approach due to its limited application.

In summary, the pangenome analysis of *Vibrio europaeus* reveals a higher proportion of core genes compared with other pathogenic vibrios, suggesting a conserved virulence arsenal and ecological specialization to shellfish hosts. Besides, the presence of multiple secretion systems highlights its pathogenic potential, demonstrated in juveniles infection assays. Finally, the high number anti-phage defense systems encoded in the accessory genome explains almost all the variability of the species. This study presents the first characterization of the pangenome of the bivalve pathogen *Vibrio europaeus*, contributing to increase the knowledge about the genomics and ecology of the species.

## FUNDING & ACKNOWLEDGMENTS

This publication derived from the R&D project PID2020-120503RA-I00 funded by MICIU/AEI/10.13039/501100011033. This study is set within the framework of the « Laboratoire d’Excellence (LabEx) » TULIP (ANR-10-LABX-41). We thank the French National Reference Laboratory for mollusc diseases (NRL, La Tremblade) for providing some of the bacterial strains used in this study.

