## Supplemental material for "Deciphering the pangenome of the shellfish pathogen *Vibrio europaeus*: Evolutionary history and functional impact of core and accessory genes in aquaculture"

**Supplementary Material**

| **Strain** | **Assembly*^1^** | **CDS** | **Accession number** |
| --- | --- | --- | --- |
| EX1 | comp (4) | 4936 | CP180205-CP180207 |
| PP-654 | cont (38) | 4921 | JAPFJT000000000.1 |
| PP-660 | cont (34) | 4891 | JAPFJS000000000.1 |
| PP-635 | cont (38) | 5014 | JAPFJR000000000.1 |
| CECT8136T | comp (4) | 4936 | LUAX00000000.1 |
| CECT8427 | comp (4) | 4941 | JAPFJQ000000000.1 |
| CECT8426 | comp (3)*^2^ | 4935 | GCA_015654285.1 |
| 07/038 2T2 | cont (44) | 5008 | JAPFJP000000000.1 |
| 07/108 T1 | cont (43) | 4887 | JAPFJO000000000.1 |
| 07/110 T1 | cont (43) | 4880 | JAPFJN000000000.1 |
| 07/112 T1 | cont (36) | 4878 | JAPFJM000000000.1 |
| 07/115 T2 | cont (47) | 4856 | JAPFJL000000000.1 |
| 07/116 T1 | cont (37) | 4851 | JAPFJK000000000.1 |
| 07/117 T1 | cont (37) | 4908 | JAPFJJ000000000.1 |
| 07/120 T1 | cont (41) | 4998 | JAPFJI000000000.1 |
| 07/121 1T1 | cont (39) | 4883 | JAPFJH000000000.1 |
| PP2-843 | comp (4) | 5086 | GCA_028447005.1 |
| PP2-978 | cont (32) | 5015 | JAPFJF000000000.1 |
| 2909 | cont (34) | 5019 | JAPFJE000000000.1 |
| 2895 | cont (33) | 4884 | JAPFJD000000000.1 |
| 2930 | cont (36) | 4952 | JAPFJC000000000.1 |
| 2951 | cont (37) | 4944 | JAPFJB000000000.1 |
| 2945 | cont (33) | 4844 | JAPFJA000000000.1 |
| 2967 | cont (33) | 5015 | JAPFIZ000000000.1 |
| 2968 | cont (34) | 4940 | JAPFIY000000000.1 |
| 2969 | cont (37) | 4941 | JAPFIX000000000.1 |
| 2971 | cont (35) | 4943 | JAPFIW000000000.1 |
| 2974 | cont (39) | 5124 | JAPFIV000000000.1 |
| 2975 | cont (35) | 4941 | JAPFIU000000000.1 |
| 3454 | cont (45) | 5205 | JAPFIT000000000.1 |
| 3492 | cont (45) | 5203 | JAPFIS000000000.1 |
| 3610 | cont (40) | 5199 | JAPFIR000000000.1 |
| 3614 | cont (40) | 5129 | JAPFIQ000000000.1 |
| NPI1 | comp (3)*^2^ | 4811 | GCA_013154935.1 |
| 071316F | cont (85)*^2^ | 4923 | VTYH00000000.1 |
| L2 | cont (40) | 5049 | JAPFIP000000000.1 |
| L3 | cont (40) | 5011 | JAPFIO000000000.1 |
| L4 | cont (36) | 4881 | JAPFIN000000000.1 |
| L20 | cont (37) | 5011 | JAPFIM000000000.1 |

**Supplementary Table 1.** Genomic information of the V. europaeus strains (n=39) used in this study.

*^1^ Assembly level: cont (contig), compl (full resolved), number of contigs are depicted in brackets.

*^2^ Assemblies from other studies (See references in Table 1).

**Supplementary Table 2.** tRNAs found from the *V. europaeus* genomes used in this study. Strains with fully resolved genomes are marked with asterisks.

| **Strain** | **tRNAs decoding Standard 20 AA** | **tRNAs with undetermined/unknown isotypes** | **Predicted pseudogenes** | **Total tRNAs** |
| --- | --- | --- | --- | --- |
| 07/038 2T2 | 101 | 2 | 3 | 106 |
| 07/108 T1 | 100 | 1 | 0 | 101 |
| 07/110 T1 | 106 | 1 | 0 | 107 |
| 07/112 T1 | 106 | 0 | 0 | 106 |
| 07/115 T2 | 107 | 0 | 1 | 108 |
| 07/116 T1 | 107 | 0 | 0 | 107 |
| 07/117 T1 | 102 | 1 | 1 | 104 |
| 07/120 T1 | 101 | 0 | 2 | 103 |
| 07/121 1T1 | 108 | 1 | 0 | 109 |
| 071316F | 85 | 0 | 3 | 88 |
| 2895 | 104 | 2 | 0 | 106 |
| 2909 | 97 | 1 | 2 | 100 |
| 2930 | 97 | 3 | 1 | 101 |
| 2945 | 98 | 2 | 0 | 100 |
| 2951 | 100 | 3 | 0 | 103 |
| 2967 | 100 | 1 | 1 | 102 |
| 2968 | 101 | 3 | 0 | 104 |
| 2969 | 101 | 2 | 0 | 103 |
| 2971 | 105 | 3 | 1 | 109 |
| 2974 | 97 | 1 | 1 | 99 |
| 2975 | 101 | 3 | 0 | 104 |
| 3454 | 95 | 2 | 0 | 97 |
| 3492 | 102 | 3 | 0 | 105 |
| 3610 | 99 | 3 | 0 | 102 |
| 3614 | 102 | 3 | 0 | 105 |
| CECT8136T* | 116 | 2 | 0 | 118 |
| CECT8426* | 119 | 0 | 2 | 121 |
| CECT8427* | 121 | 0 | 1 | 122 |
| EX1* | 118 | 0 | 1 | 119 |
| L2 | 101 | 2 | 1 | 104 |
| L20 | 97 | 3 | 0 | 100 |
| L3 | 100 | 3 | 1 | 104 |
| L4 | 103 | 3 | 0 | 106 |
| NPI-1* | 116 | 1 | 1 | 118 |
| PP2-843* | 119 | 1 | 1 | 121 |
| PP2-978 | 102 | 1 | 2 | 105 |
| PP-635 | 100 | 3 | 0 | 103 |
| PP-654 | 100 | 2 | 0 | 102 |
| PP-660 | 100 | 3 | 0 | 103 |

**Supplementary Table 3.** rRNAs found from the *V. europaeus* genomes used in this study. Strains with fully resolved genomes are marked with asterisks.

| **Strain** | **16S rRNA** | **5S rRNA** | **23S rRNA** |
| --- | --- | --- | --- |
| 07/038 2T2 | 1 | 2 | 1 |
| 07/108 T1 | 1 | 3 | 1 |
| 07/110 T1 | 1 | 3 | 1 |
| 07/112 T1 | 1 | 3 | 1 |
| 07/115 T2 | 1 | 1 | 1 |
| 07/116 T1 | 1 | 3 | 1 |
| 07/117 T1 | 1 | 2 | 1 |
| 07/120 T1 | 1 | 1 | 1 |
| 07/121 1T1 | 1 | 2 | 1 |
| 071316F | 1 | 1 | 1 |
| 2895 | 1 | 2 | 1 |
| 2909 | 1 | 3 | 1 |
| 2930 | 1 | 2 | 1 |
| 2945 | 1 | 2 | 1 |
| 2951 | 1 | 3 | 1 |
| 2967 | 1 | 2 | 1 |
| 2968 | 1 | 3 | 1 |
| 2969 | 1 | 2 | 1 |
| 2971 | 1 | 2 | 1 |
| 2974 | 1 | 2 | 1 |
| 2975 | 1 | 4 | 2 |
| 3454 | 1 | 2 | 1 |
| 3492 | 1 | 2 | 1 |
| 3610 | 1 | 2 | 1 |
| 3614 | 1 | 2 | 1 |
| CECT8427* | 10 | 11 | 10 |
| CECT8136T* | 9 | 10 | 9 |
| CECT8426* | 10 | 11 | 10 |
| EX1* | 10 | 11 | 10 |
| L2 | 1 | 2 | 1 |
| L20 | 1 | 3 | 1 |
| L3 | 1 | 2 | 1 |
| L4 | 1 | 2 | 1 |
| NPI-1* | 9 | 10 | 9 |
| PP2-843* | 10 | 10 | 10 |
| PP2-978 | 1 | 3 | 1 |
| PP-635 | 1 | 2 | 1 |
| PP-654 | 1 | 3 | 1 |
| PP-660 | 1 | 2 | 1 |

**Supplementary Table 3.** Virulence factors identified from the *V. europaeus* pangenome.

Core virulence-related genes were indicated in bold. Virulence factor classes (VFCs) are numbered according to Fig. 4.

| **VFclass** | **Virulence factors** | **Related genes** |
| --- | --- | --- |
| Adherence (VFC 2) | Mannose-sensitive hemagglutinin (MSHA type IV pilus) | *mshA, mshB, mshC, mshE, mshG, mshH, mshI, mshJ, mshK, mshL, mshM, mshN.* |
|  | Type IV pilus | *pilA, pilB, pilC, pilD.* |
|  | Curli fibers(*Escherichia*) | ***csgG*** |
|  | LPS-O-antigen (P. aeruginosa)(*Pseudomonas*) | *tviB, tviB*(2), *hisF2*. |
|  | Tap type IV pili(*Aeromonas*) | *tapQ* |
|  | The tad locus(*Haemophilus*) | ***tadA****,* ***tadA*(2)** |
|  | Type IV pili(*Yersinia*) | *pilW* |
| Antiphagocytosis (VFC 3) | Capsular polysaccharide | *cpsA, cpsC, rmlA, rmlB, rmlC, rmlD, wbfT, wbfU, wbfV/wcvB, wbfY.* |
|  | Capsule (*Klebsiella*) | *uge* |
| Chemotaxis and motility (VFC 5) | Flagella | ***cheA****,* ***cheA*(2)**, ***cheB****,* ***cheR****,* ***cheV****,* ***cheW****,* ***cheY****,* ***cheZ****,* ***filM****,* ***flaA****,* ***flaB****,* ***flaC****,* ***flaD****,* ***flaE****,* ***flaG****,* ***flaI****,* ***flgA****,* ***flgB****,* ***flgC****,* ***flgD****,* ***flgE****,* ***flgF****,* ***flgG****,* ***flgH****,* ***flgI****,* ***flgJ****,* ***flgK****,* ***flgL****,* ***flgM****,* ***flgN****,* ***flhA****,* ***flhB****,* ***flhF****,* ***flhG****,* ***fliA****,* ***fliD****,* ***fliE****,* ***fliF****,* ***fliG****,* ***fliH****,* ***fliI****,* ***fliJ****, fliK,* ***fliL****,* ***fliN****,* ***fliO****,* ***fliP****,* ***fliQ****,* ***fliR****,* ***fliS****,* ***flrA****,* ***flrB****,* ***flrC****,* ***motA****,* ***motB****,* ***motX****,* ***motY****.* |
| Enzyme (VFC 7) | Metalloproteinase | ***hap/vvp****,* ***prtV*** |
|  | Collagenase | ***colA****,* ***colP*** |
| Iron uptake (VFC 10) | Enterobactin receptors | ***vctA*** |
|  | Heme receptors | ***hutA*** |
|  | Periplasmic binding protein-dependent ABC transport systems | *vctC, vctD, vctG, vctP, vctP*(2) |
|  | Pyoverdine(*Pseudomonas*) | *pvdY* |
| Quorum sensing (VFC 13) | Autoinducer-2 | ***luxS*** |
|  | Cholerae autoinducer-1 | ***cqsA*** |
| Secretion system (VFC 14) | EPS T2SS | ***epsC, epsE, epsF, epsG, epsH, epsI, epsJ, epsK, epsL, epsM, gspD*** |
|  | T3SS1 secreted effectors | Undetermined, *vopQ* |
|  | T3SS1 | *sycN, tyeA, vcrD, vcrG, vcrH, vcrR, virF, virG, vopB, vopN, vscA, vscB, vscC, vscD, vscF, vscG, vscI, vscJ, vscK, vscL, vscN, vscO, vscQ, vscR, vscS, vscT, vscU, vscX, vscY, vxsC.* |
|  | VAS effector proteins | *hcp-2, hcp-2*(2), *vgrG*-2*, vgrG*-2*(2), vgrG-*2(3), *vgrG*-3 |
|  | VAS T6SS secretion system | ***vasA****,* ***vasB****,* ***vasC****,* ***vasD****,* ***vasE****,* ***vasF****,* ***vasG***, ***vasH***, ***vasJ***, ***vasK***, ***tssE***, ***impC*/*vipB***, ***ipmB*/*vipA***, *impA*, ***impA*(2)**, *hcp*, *hcp*(2), ***hcp*(3)***, impB/vipA,* ***impB/vipA*(2)**, *impC/vipB, i****mpC/vipB*(2)**, ***impC/vipB*(3)**, *tssE,* ***tssE*(2)**, *impG/vasA,* ***impG/vasA*(2)**, *impH/vasB,* ***impH/vasB*(2)**, *tssH,* ***tssH*(2)**, *impI/vasC,* ***impI/vasC*(2)**, *vasD,* ***vasD*(2)**, *impJ/vasE,* ***impJ/vasE*(2)**, *impK/vasF/ompA/motB,* ***impK/vasF/ompA/motB* (2)**, *icmF/vasK,* ***icmF/vasK*(2)**, PAAR-repeat_protein, **PAAR-repeat_protein(2)**. |
|  | T3SS(*Aeromonas*) | *ati1* |
| Toxin (VFC 15) | RTX toxin | *rtxB, rtxD* |
|  | *V.cholerae* cytolysin | *hlyA* |
|  | *V.cholerae* cytolysin | *hlyA* |
|  | *V.cholerae* cytolysin | *hlyA* |
|  | *V.cholerae* cytolysin | *hlyA* |
|  | *V.cholerae* cytolysin | *hlyA* |
| Acid resistance (VFC 1) | Urease(*Helicobacter*) | ***ureB****,* ***ureG*** |
| Biofilm formation (VFC 4) | AdeFGH efflux pump/transport autoinducer(*Acinetobacter*) | ***adeG*** |
| Endotoxin (VFC 6) | LOS(*Haemophilus*) | *kdsA, kpsF* |
|  | LPS(*Bordetella*) | *bplA* |
| Glycosylation system (VFC 8) | O-linked flagellar glycosylation(*Campylobacter*) | *pseB* |
| Immune evasion (VFC 9) | Capsule(*Neisseria*) | *ctrD* |
|  | Capsule(*Acinetobacter*) | *pseI, pseF, pseC, wbpD, tviB(3), pseB(2), vipB/tviC, wbpP* |
| Nutritional factors (VFC 11) | Allantoin utilization(*Klebsiella*) | *allA* |
| Others (VFC 12) | O-antigen(*Yersinia*) | *manA, wcaG* |

**Supplementary Table 4.** Differences in the non-core virulence factors encoded by the strains belonging to the same phylogenetic cluster.

Strains encoding a certain gene are indicated in bold.

| **Sub-cluster Ia** | immune evasion gene *ctrD* (**L4**) |
| --- | --- |
| **Sub-cluster Ib** | VAS effector protein coding genes *vgrG-2* (**2967, 2974** and **2909**) and *vgrG-2_3* (**PP2-978**); toxins coding genes *cysC1* (**2967, 2974** and **PP2-978**) or *rtxE* (**2909**); allantoin utilization gene *allA* (**2967**).  **L2:** VAS effector proteins of the T6SS (*hcp2_2, vgrG-2, vgrG-2_3* and *vgrG-3*), toxins (*rtxB, rtxD, rtxE, hlyA* and *cysC1*), endotoxins (*kdsA* and *kpsF*), immune evasion (*ctrD*) and allantoin utilization (*allA*). |
| **Subcluster IIa** | No differences |
| **Subcluster IIb** | type 4 pili *tapQ* and the toxin *cysC1* (**07/120 T1**); *rtxE* and *allA* (**CECT8426**). |
| **Cluster III** | type IV pili *pilW* and VAS effectors *hcp-2_2* and *vgrG2_3* (**07/117 T1**). |

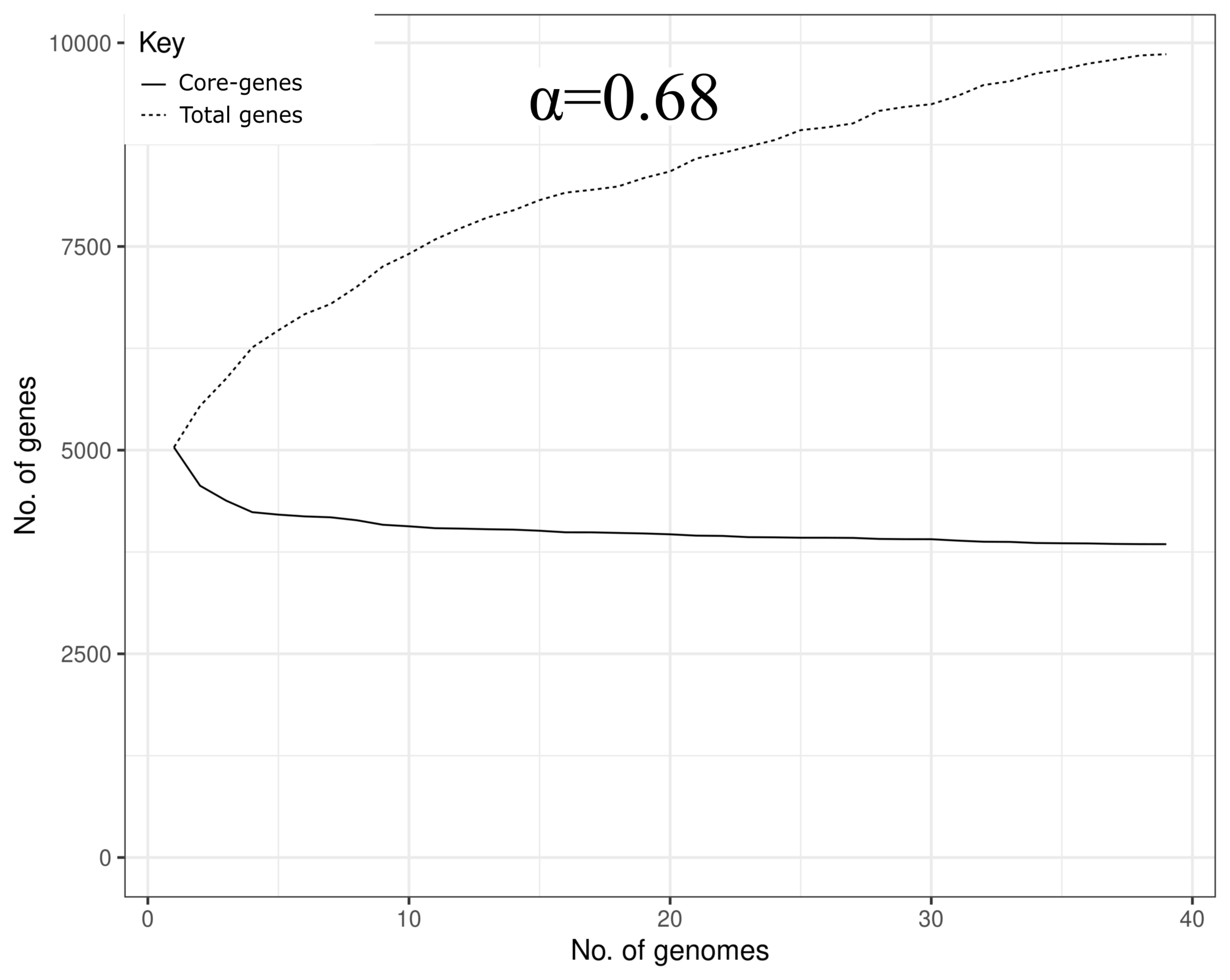

**Supplementary Figure 1.** Relation between the number of genes (core and total) and the number of genomes. α coefficient (value=0.68) of Heap's Law for conserved curve is shown.

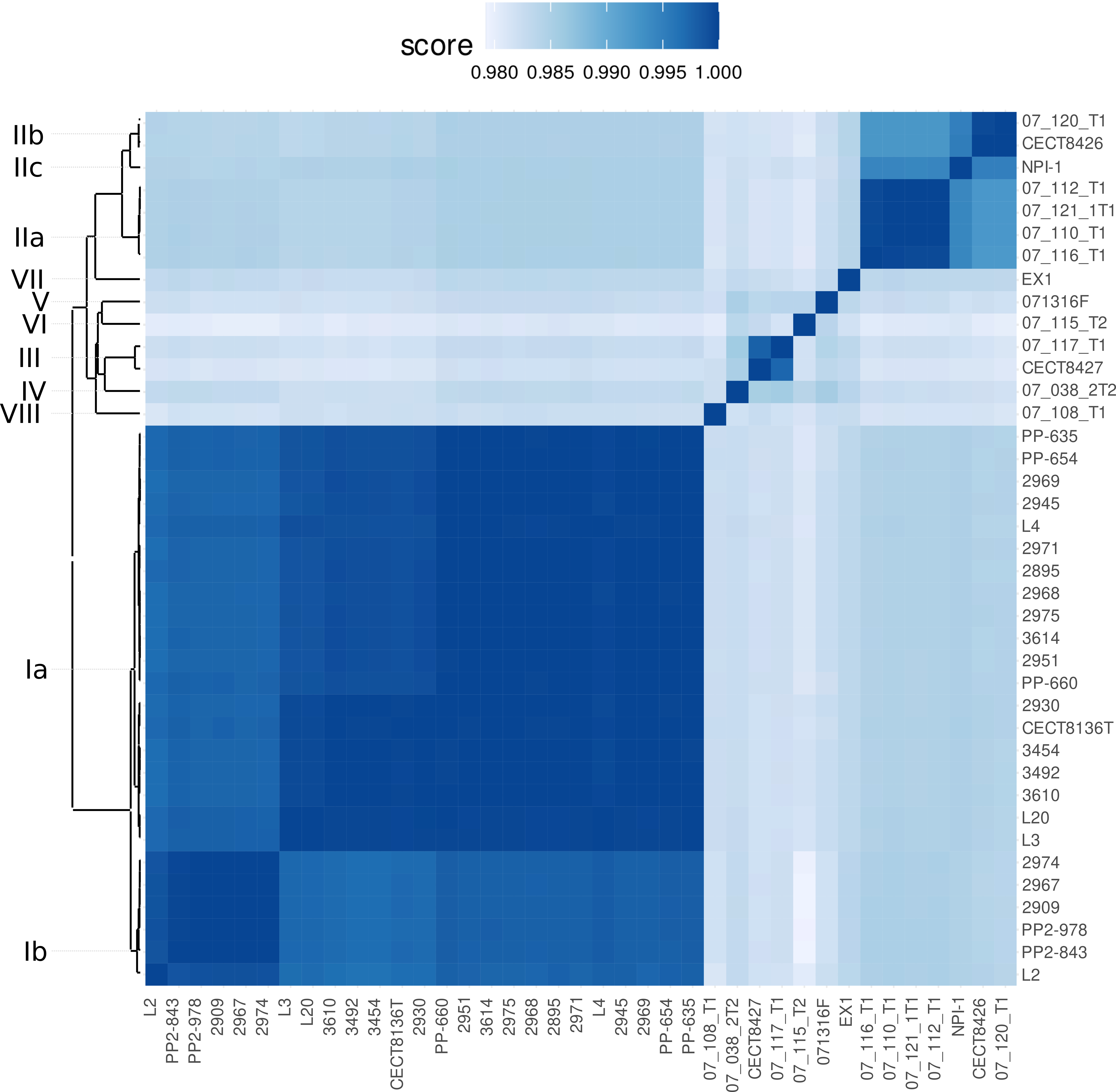
**Supplementary Figure 2.** ANI heatmap of V. europaeus strains with hierarchical clustering.

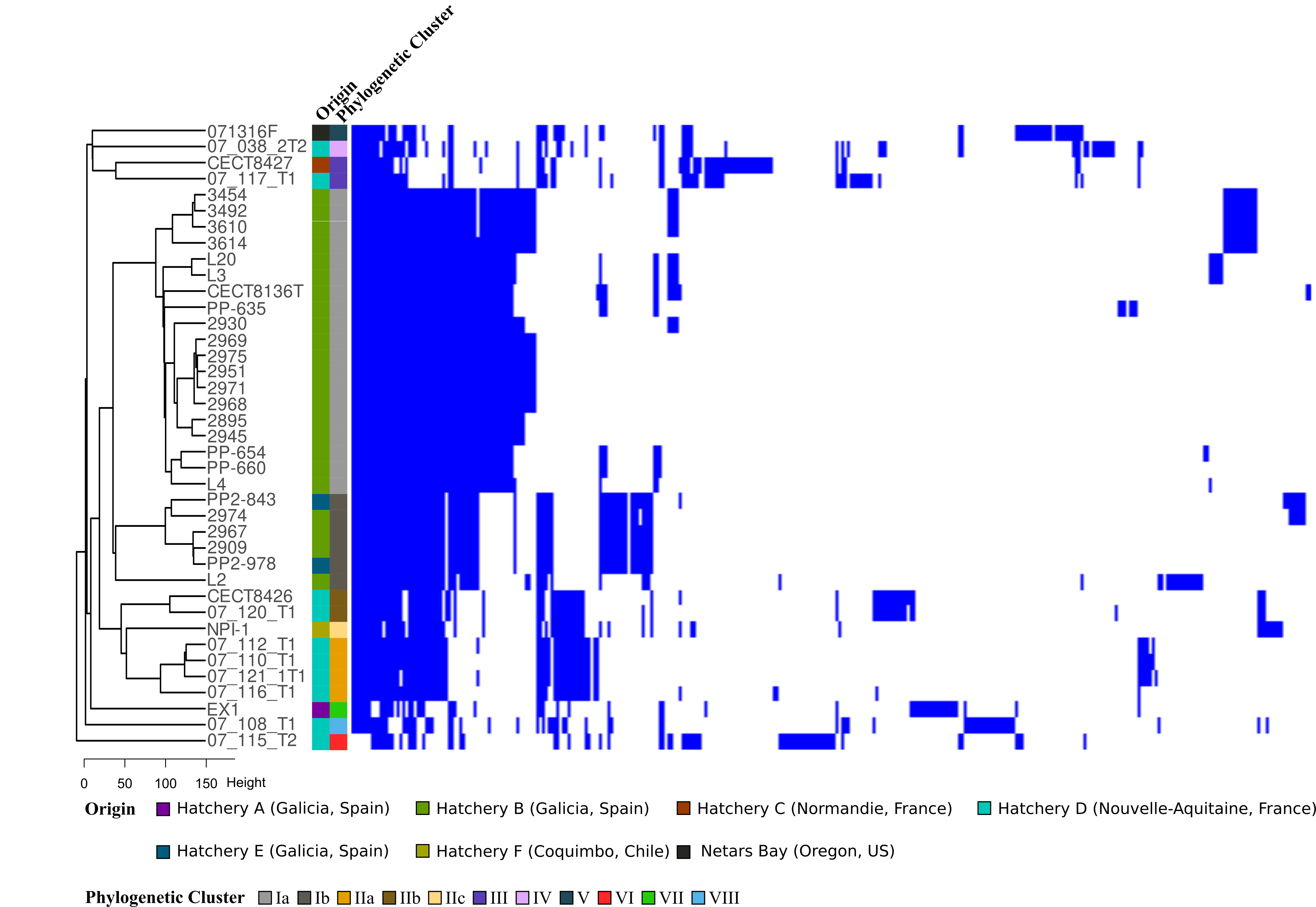

**Supplementary Figure 3.** Presence/absence matrix showing the *V. europaeus* accessory genes. Strains are sorted according to the dendrogram plotted to the left.

A) T6SS1

B) T6SS2

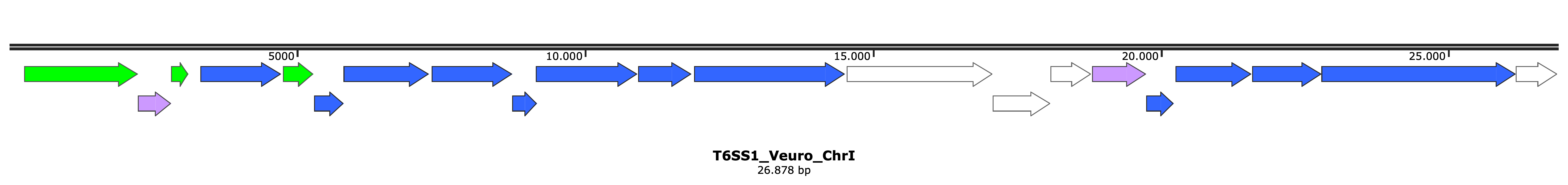

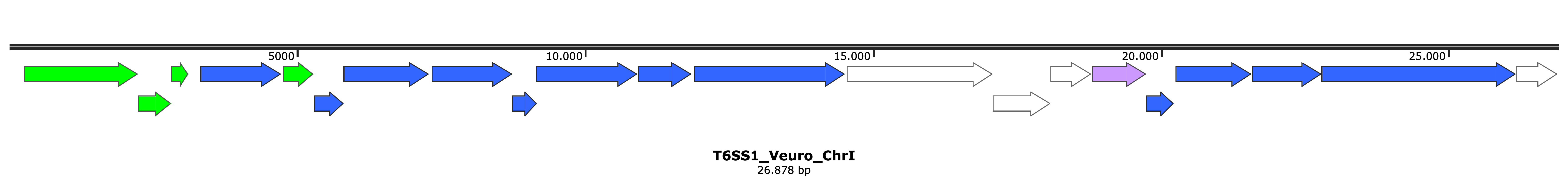

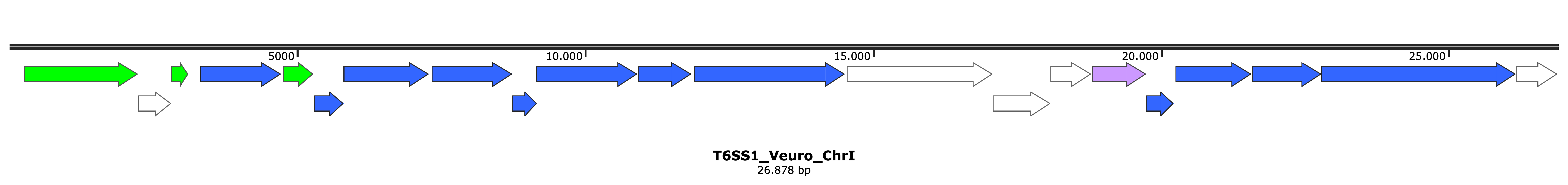

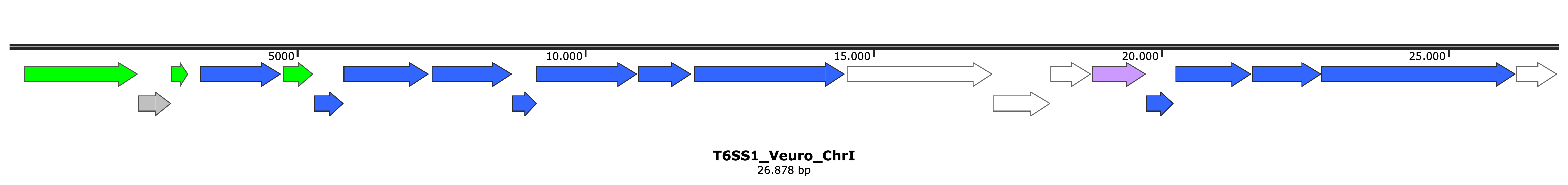

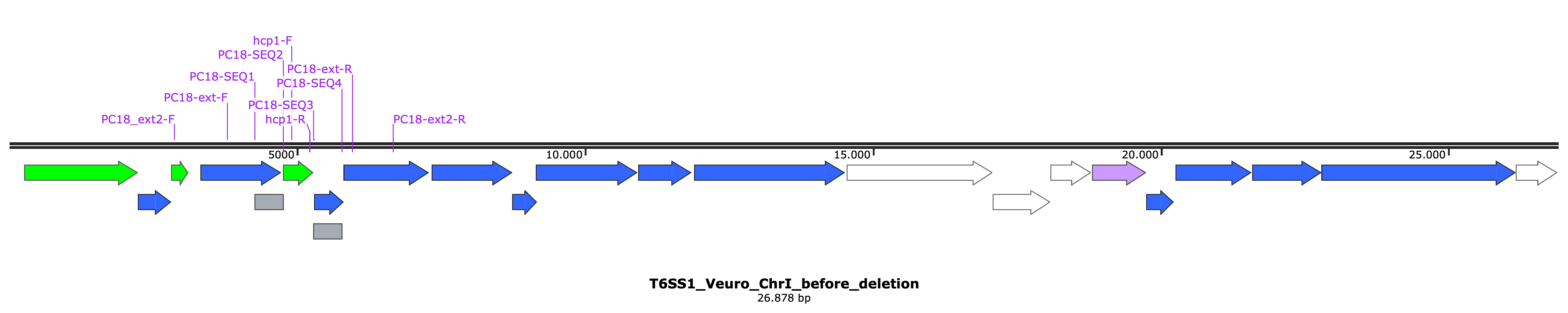

core

tube-spike

other

accessory

pseudogene

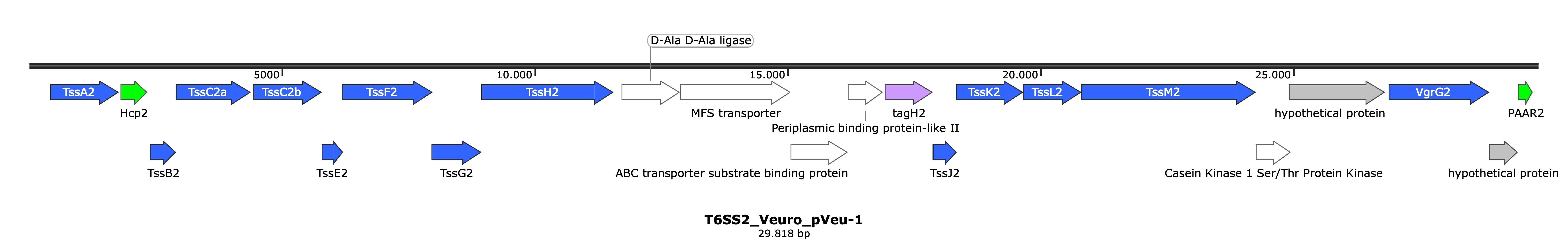

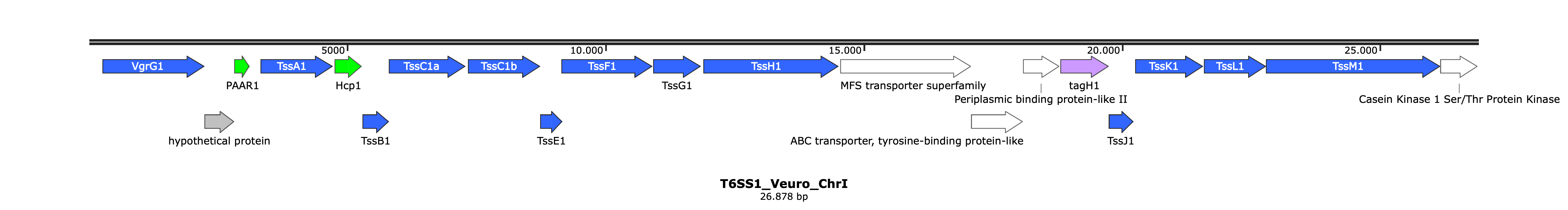

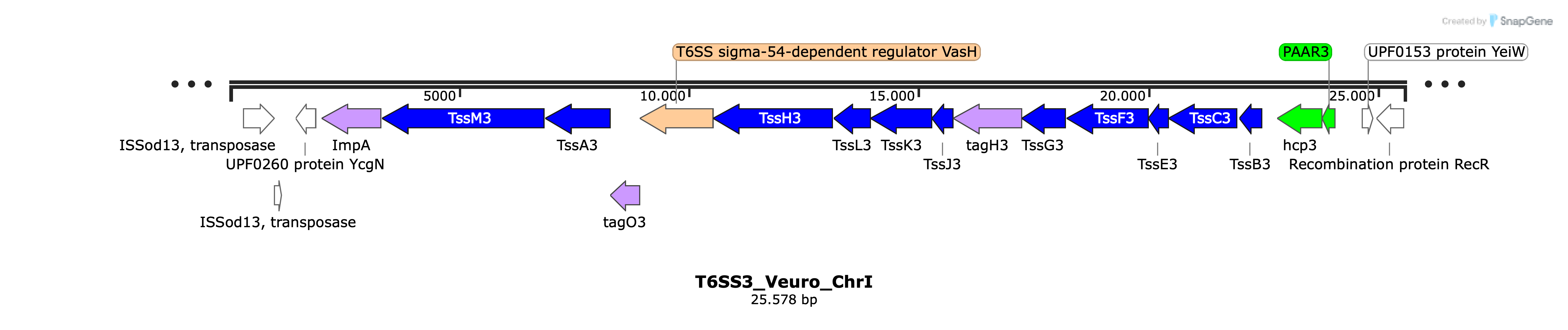

C) T6SS3

**Supplementary Figure 4.** T6SS (T6SS1, A; T6SS2, B; T6SS3, C) gene clusters found in *V. europaeus* genomes. T6SS1 and T6SS3 were assigned to the core-genome. T6SS genes are denoted by arrows indicated the predicted direction of the transcription. Encoded proteins or domains are denoted on the genes.

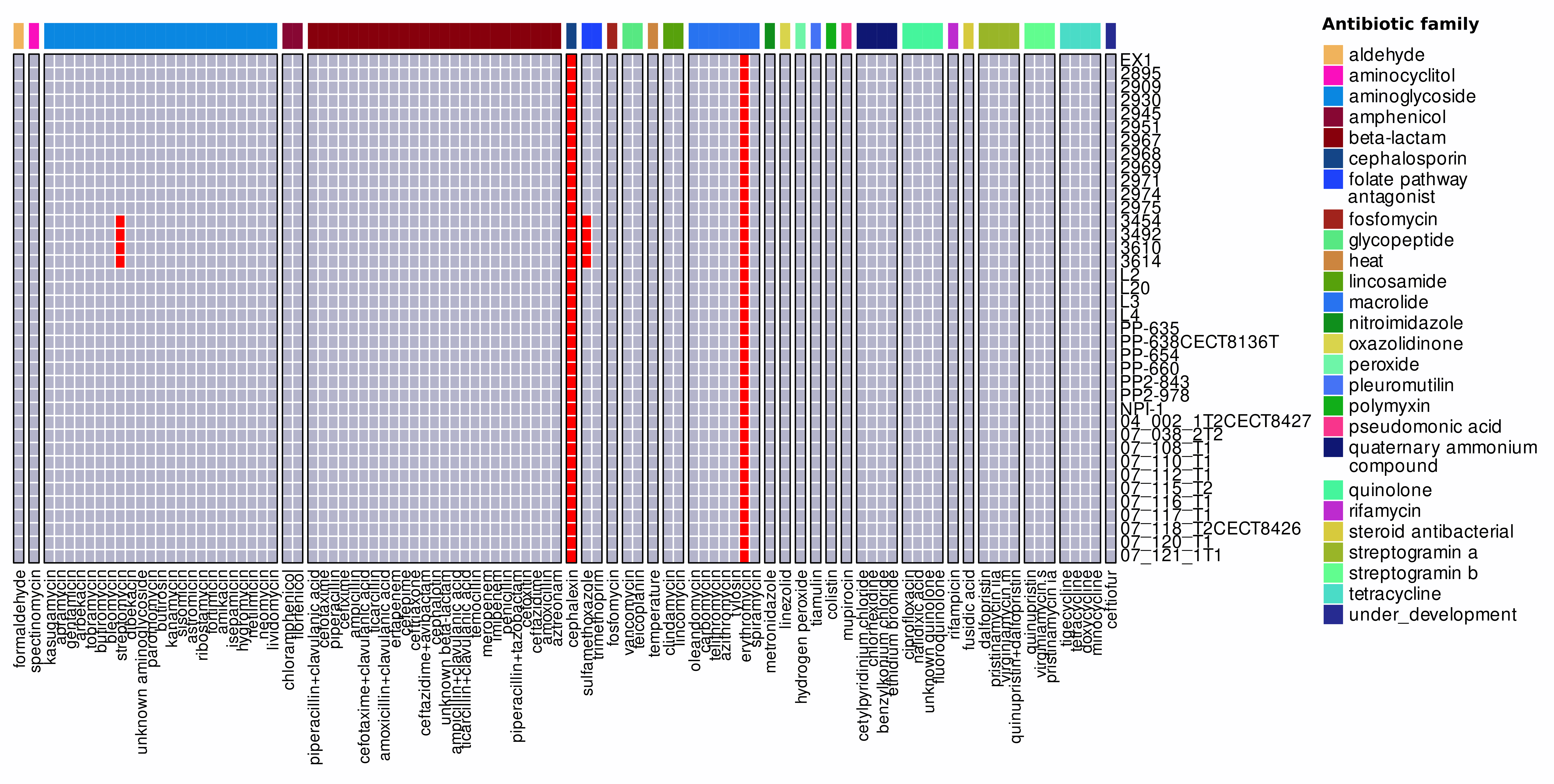

**Supplementary Figure 5.** Antibiotic resistance phenotypes of the *V. europaeus* strains.

Antibiotics are depicted by columns and grouped by antibiotic families as indicated in the right legend
